# Decoding m^6^Am by simultaneous transcription-start mapping and methylation quantification

**DOI:** 10.1101/2024.10.16.618717

**Authors:** Jianheng Fox Liu, Ben R. Hawley, Luke S. Nicholson, Samie R. Jaffrey

## Abstract

*N*^6^,2’-*O*-dimethyladenosine (m^6^Am) is a modified nucleotide located at the first transcribed position in mRNA and snRNA that is essential for diverse physiological processes. m^6^Am mapping methods assume each gene uses a single start nucleotide. However, gene transcription usually involves multiple start sites, generating numerous 5’ isoforms. Thus, gene levels annotations cannot capture the diversity of m^6^Am modification in the transcriptome. Here we describe CROWN-seq, which simultaneously identifies transcription-start nucleotides and quantifies m^6^Am stoichiometry for each 5’ isoform that initiates with adenosine. Using CROWN-seq, we map the m^6^Am landscape in nine human cell lines. Our findings reveal that m^6^Am is nearly always a high stoichiometry modification, with only a small subset of cellular mRNAs showing lower m^6^Am stoichiometry. We find that m^6^Am is associated with increased transcript expression and provide evidence that m^6^Am may be linked to transcription initiation associated with specific promoter sequences and initiation mechanisms. These data suggest a potential new function for m^6^Am in influencing transcription.

## INTRODUCTION

m^6^Am (*N*^6^,2’-*O*-dimethyladenosine) is the most common modified nucleotide in mRNA. m^6^Am is found specifically at the first transcribed position of mRNAs, termed the transcription-start nucleotide (TSN), which reflects the transcription-start site (TSS) in DNA. During transcription, the TSN typically acquires a 2’-*O*-methyl modification^1,2^, which is deposited by CMTR1 (Cap-specific mRNA nucleoside-2’-*O*-methyltransferase 1)^3^. In the case of mRNAs that use adenosine as the TSN, the initial 2’-*O*-methyladenosine (Am) can be further methylated on the *N*^6^ position of the adenine nucleobase to form m^6^Am by PCIF1 (Phosphorylated CTD Interacting Factor 1)^4^.

Studies using PCIF1 depletion (i.e., m^6^Am depletion) have revealed that m^6^Am has important roles in cell physiology. In normal cells, PCIF1 depletion does not appear to affect cell growth or viability^4^. However, in oxidative stress conditions, PCIF1 deficiency leads to impaired cell growth^4^. In cancer cells, PCIF1 depletion markedly enhances cell death during anti-PD1 therapy^5^. During viral infection, PCIF1 depletion results in increased HIV replication^6^, impaired SARS-Cov-2 infection^7^, and increased VSV immunogenicity^8^. These studies indicate that m^6^Am has important roles in diverse cellular contexts.

A major goal has been to identify and characterize the m^6^Am- and Am-containing mRNAs. Initial chromatographic studies in the 1970s demonstrated that cellular mRNAs can exist in m^6^Am and Am forms, with the Am form being more predominant^9^. To map m^6^Am modified genes, several antibody-based methods were developed, including miCLIP^10^, m^6^Am-seq^11^, m6ACE-seq^12^, and m6Am-exo-seq^13^. These methods can identify m^6^Am sites (miCLIP and m6ACE-seq), m^6^Am peaks (m^6^Am-seq), or m^6^Am containing genes (m^6^Am-exo-seq). Am genes were identified when m^6^Am was not detected but the reported TSN in publicly available datasets was A. The remaining genes were annotated as Gm, Cm, or Um based on public TSS annotations.

However, despite these transcriptome-wide m^6^Am maps, the effect of m^6^Am on mRNA is unclear. By examining m^6^Am genes, along with the change in mRNA stability and translation, small and inconsistent effects have been observed from different labs^4,13–15^.

We considered the possibility that the difficulty in establishing m^6^Am function may be due to flaws in the way that genes are designated as m^6^Am genes. Previous m^6^Am mapping studies treated m^6^Am like other modified nucleotides, which are internal. In studies of internal nucleotides, such as m^6^A, isoform diversity is generally not considered since these different isoforms rarely impact the detection of the nucleotide. In contrast, m^6^Am is highly affected by isoform diversity since it is located at the 5’ end. Most genes generate multiple transcript 5’ isoforms that each use a different TSSs^16^. However, previous m^6^Am mapping studies assumed that each gene has a single TSN, whose identity was based on existing gene annotations. Therefore, existing m^6^Am maps that assign a specific start nucleotide to each gene cannot be accurate, since most genes produce a range of transcripts, with possibly multiple start nucleotides. For this reason, m^6^Am mapping and functional studies of m^6^Am need to be performed in a way that considers the 5’ isoform diversity of most genes.

Another concern is that the existing m^6^Am mapping studies designated each gene as either m^6^Am or Am. However, it is possible that m^6^Am levels can be variable, with only a fraction of transcripts containing m^6^Am and the remainder containing Am. Stoichiometric maps of m^6^Am can potentially reveal the degree to which a transcript would be influenced by m^6^Am-dependent pathways.

Quantitative m^6^Am mapping is especially important for small nuclear RNAs (snRNAs), which also contain m^6^Am at their TSN. Initial biochemical characterization of snRNAs revealed that the first nucleotide was Am^17^, but subsequent studies showed that nearly half of all snRNAs are initially methylated to m^6^Am, and then demethylated by FTO to Am^10,12^. Thus, m^6^Am is a transient intermediate in snRNA biogenesis. Notably, m^6^Am levels can be highly regulated in snRNAs^10^, however, current m^6^Am mapping methods are unable to quantify changes in m^6^Am stoichiometry.

To understand the transcriptome-wide distribution of m^6^Am, we developed CROWN-seq (**<u>C</u>**onversion **<u>R</u>**esistance detection **<u>O</u>**n **<u>W</u>**hole-transcriptomic transcription-start ***<u>N</u>***^6^,2’-*O*-dimethyladenosine by **<u>seq</u>**uencing), an antibody-free quantitative m^6^Am mapping method. Using CROWN-seq, we define the overall repertoire of 5’ isoforms for each gene, and the specific isoforms that use m^6^Am as the TSN across nine different cell lines. We find that annotations of genes based on a single start nucleotide do not capture the diversity of 5’ transcript isoforms for most genes. Instead, m^6^Am is more accurately assessed for each 5’ isoform separately. Nearly all A-initiated transcript isoforms have very high m^6^Am stoichiometry, and that transcripts containing Am as the TSN are relatively rare. Transcript isoforms that contain m^6^Am are more highly expressed, and loss of m^6^Am due to depletion of PCIF1 leads to reduced expression of transcript isoforms containing m^6^Am. However, we find that this effect is not due to decreased mRNA stability. Instead, the depletion of PCIF1 affects transcripts based on upstream core promoter elements. Our data suggest that transcription mechanisms that utilize specific core promoter sequences achieve high expression, which might be linked to a transcription-promoting effect of m^6^Am. Overall, our quantitative transcriptome-wide transcription-start nucleotide m^6^Am maps reveal a markedly distinct m^6^Am profile than previously measured, show that m^6^Am is the predominant modified nucleotide relative to Am in mRNA, and suggest roles of m^6^Am in transcription.

## RESULTS

### ReCappable-seq reveals high transcript isoform diversity at the 5’ end

In previous m^6^Am mapping studies, genes were annotated to be m^6^Am, Am, Gm, Cm, or Um ^4,10,13,14^, based on the assumption that each gene has one major transcription-start nucleotide. To determine how often genes are characterized by a single major TSS we used ReCappable-seq^18^, a method for quantitative measurement of transcription-start sites. ReCappable-seq is similar to traditional TSS-seq methods which involve ligation of an oligonucleotide to the 5’ end of mRNAs^19^, thus precisely marking the TSN. However, ReCappable-seq adds an enrichment step for capped mRNA fragments to significantly reduce background signals from internal sites that are derived from RNA cleavage. Thus, ReCappable-seq provides a highly sensitive and precise mapping of TSNs at single-nucleotide resolution (**Note S1**, “The comparison of transcription-start mapping methods”).

By analyzing ReCappable-seq data in HEK293T cells, we found that protein-coding genes tend to have multiple TSNs. Among the 9,199 genes analyzed, we identified 87,624 TSNs (see **Methods**). On average, a gene uses 9.5±9 (mean and s.d., hereafter) TSNs (**Figure 1A**). Only ∼9% of genes contain a single TSN (**Figure 1A**). Thus, most genes cannot be characterized by a single start nucleotide.

**Figure 1.**
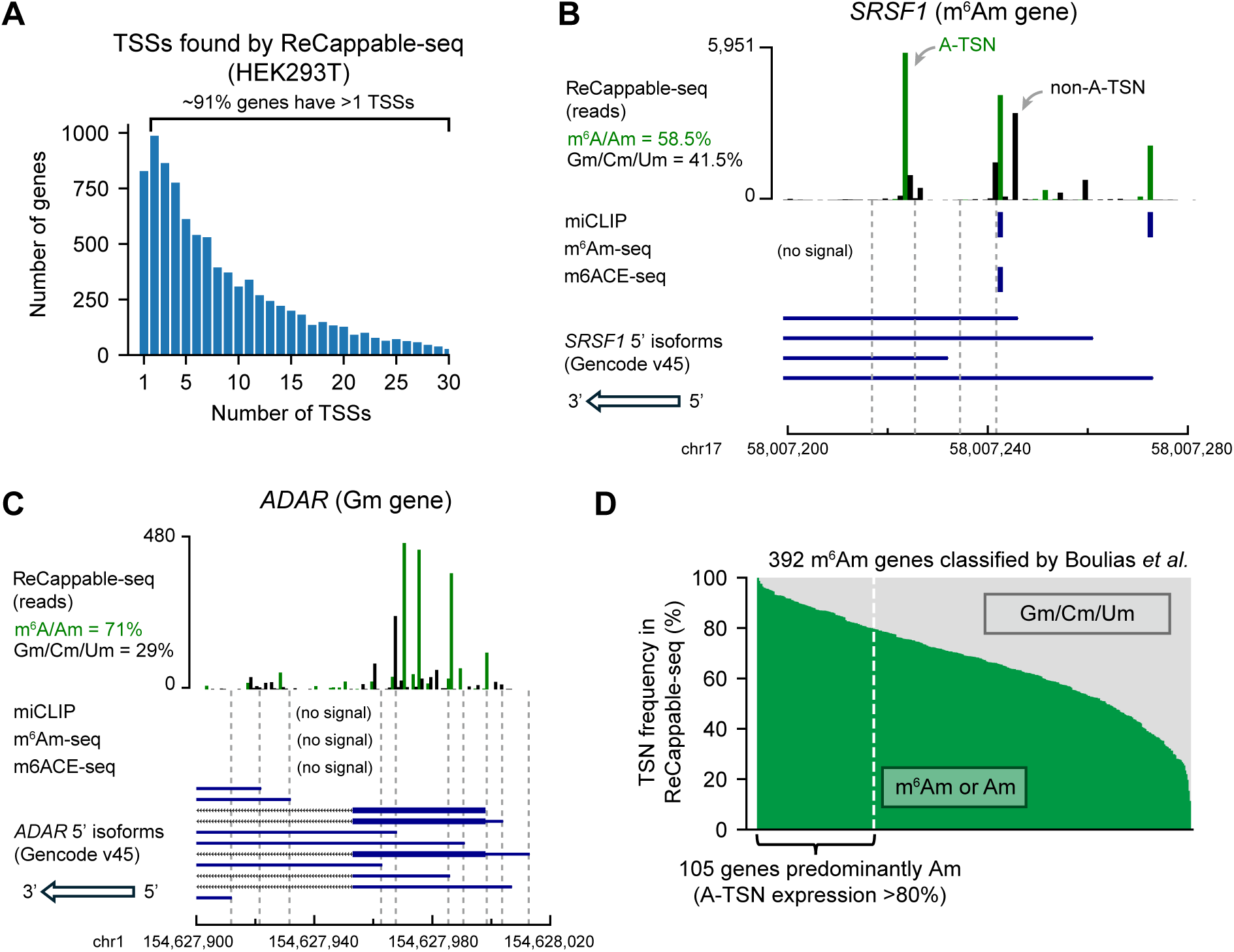
Many m^6^Am genes are mistakenly annotated. **(A)** Genes tend to have multiple TSSs. Shown is a histogram displaying the number of TSSs per protein-coding gene in HEK293T cells. TSSs (87,624 TSSs from 9,199 genes) were mapped using ReCappable-seq. These TSSs have expression levels ≥1 TPM (transcription-start nucleotide per million). **(B)** and **(C)** Examples of genes that were mistakenly classified by previous studies^14^. *SRSF1* **(B)** was previously designated as m^6^Am because of the miCLIP signals overlapping with A-TSSs. However, based on ReCappable-seq, ∼41.5% of the reads are mapped to non-A-TSSs in *SRSF1*. Notably, one of the most expressed m^6^Am A-TSS (chr17:58,007,228) was mistakenly considered as internal m^6^A because this position was not previously annotated as a TSS^44^. *ADAR* **(C)** was previously classified as Gm. There is no m^6^Am signal based on miCLIP^14^, m^6^Am-seq^11^, or m6ACE-seq^12^ mapped to *ADAR*. However, ∼71% of the transcripts from *ADAR* are A- initiated. **(D)** Previously classified m^6^Am genes express considerable levels of non-A-initiated transcripts. Each column represents a gene previously classified as m^6^Am gene by miCLIP^14^. For each gene, the percentage of transcript isoforms starting with m^6^Am/Am (in green) or Gm/Cm/Um (in gray) are shown. The percentage was calculated by weighting each transcript isoform by its expression level. The TSN frequencies were obtained using ReCappable-seq in HEK293T cells.

As an example, *SRSF1*, which was previously classified as a m^6^Am gene, has ∼37 different 5’ isoforms in HEK293T cells, of which ∼41.5% do not use an A-TSN (**Figure 1B**). As another example, *ADAR* was previously classified as a Gm gene, but ∼71.0% of transcripts use A-TSNs (**Figure 1C**). These observations are not artifacts of ReCappable-seq because similar results were also found using other TSN mapping methods (**Figure S1A, S1B**).

Conceivably m^6^Am genes produce multiple 5’ isoforms, but the isoforms predominantly use A- TSNs. If this were the case, then the gene could indeed be considered an m^6^Am gene if all the A-TSNs were methylated to m^6^Am. We considered a gene to be predominantly composed of A- TSNs if >80% of transcripts start with A. Using this criterion, we found that only ∼24% of m^6^Am genes determined by miCLIP^14^ are primarily composed of A-TSNs (**Figure 1D**). Similar observations were also found in other m^6^Am mapping methods^4,13^ (**Figure S1C, D**).

Our ReCappable-seq analysis also suggested that previous m^6^Am mapping methods may not have detected the diversity of m^6^Am in the transcriptome. ReCappable-seq identified many more A-TSNs than the total number of previously mapped m^6^Am sites. For example, in both *SRSF1* and *ADAR*, many A-TSNs are seen using ReCappable-seq, however, m^6^Am signals were only found at a few of these A-TSNs by either miCLIP^14^, m^6^Am-seq^11^, or m6ACE-seq^12^ (**Figure 1B, C**). This might suggest that only a few A-TSNs are m^6^Am modified. However, it is also possible that the antibody-based mapping methods do not have the resolution or sensitivity to distinguish between m^6^Am at different 5’ isoforms. Notably, previous m^6^Am mapping studies exhibited very low overlap with each other. miCLIP^14^, m^6^Am-seq^11^ and m6ACE-seq^12^ together identified 7,480 m^6^Am sites in HEK293T cells (**Figure S1E** and **Table S1**). Among these sites, only 1.1% (84) are found in all three methods and 9.7% (728) are found in at least two studies (**Figure S1E** and **Table S1**). Taken together, these data demonstrate a variety of concerns about existing m^6^Am mapping studies.

### CROWN-seq integrates TSN mapping and m^6^Am quantification

To understand the distribution of m^6^Am in the transcriptome, we sought to develop a method to identify the entire repertoire of TSNs among all the 5’ transcript isoforms for each gene. In this way, we can identify the specific 5’ isoforms for each gene that contain m^6^Am. Additionally, we wanted a quantitative method rather than the qualitative assessment provided by previous antibody-based methods. Recently, chemical methods using sodium nitrite were developed for m^6^A analysis^20–22^. This method identifies m^6^A by chemically deaminating (“converting”) unmethylated A’s into inosines (I’s), while leaving m^6^A’s intact. During sequencing, the A-to-I conversions are readily detected because I’s are reverse transcribed into G’s. This approach leads to precise and robust m^6^A quantification^20^. Because of the chemical similarity between m^6^Am and Am, we explored the potential use of sodium nitrite conversion to map and quantify m^6^Am.

We first asked if Am is susceptible to deamination by sodium nitrite. To test this, we performed sodium nitrite conversion on a m^7^G-ppp-Am-initiated transcript (see **Methods**). We applied the sodium nitrite conversion protocol used in GLORI, which includes glyoxal treatment to prevent modification of guanosine residues^20^. After sodium nitrite treatment, the RNA was reverse transcribed and sequenced. The conversion rate of Am was quantified by counting A or G reads at the first nucleotide position. In this assay, Am was completely converted (**Figure S2A**), indicating that sodium nitrite efficiently converts Am and thus can be used for m^6^Am quantification.

We considered the possibility that GLORI data^20^ could be mined to measure m^6^Am stoichiometry at previously mapped m^6^Am sites^14^. We noticed that many A’s at these TSNs were highly converted in GLORI (**Figure S2B**), suggesting prevalent Am. This is inconsistent with mass spectrometry analysis of mRNA cap structures from us^1^ and others^2,4^, which has suggested that m^6^Am is very prevalent while Am is relatively rare in mRNA. A potential cause of the high level of Am at TSNs predicted by GLORI could be the extensive RNA fragmentation that occurs with sodium nitrite treatment. RNA fragments that have 5’ ends that align to the TSNs of overlapping transcript isoforms can confound the measurement of m^6^Am stoichiometry (**Figure S2C, S2D**). Thus, GLORI cannot distinguish between true TSNs and internal bases that are found at the 5’ end of RNA fragments. To overcome this limitation, we sought to develop a method that selectively analyzes A-TSNs and thereby removes the confounding effect of overlapping transcripts.

We developed CROWN-seq, which selectively analyzes TSNs throughout the transcriptome (**Figure 2A**). In this method, Am residues in mRNA are converted to Im using sodium nitrite. Next, we specifically isolate the 5’ ends of mRNA by replacing the m^7^G cap with a desthiobiotin affinity tag using a decapping-and-recapping strategy^18^. By enriching the m^7^G-proximal sequence, we can simultaneously sequence the TSN of all transcripts, including both m^6^Am and non-m^6^Am TSNs. This is conceptually different from existing m^6^Am mapping methods which only examine the m^6^Am transcripts. For A-TSNs, m^6^Am stoichiometry can be quantified by counting the number of A reads (reflecting m^6^Am) or G reads (reflecting Am). In this way, we not only obtain TSN locations but also m^6^Am stoichiometry in the same RNA molecule.

**Figure 2.**
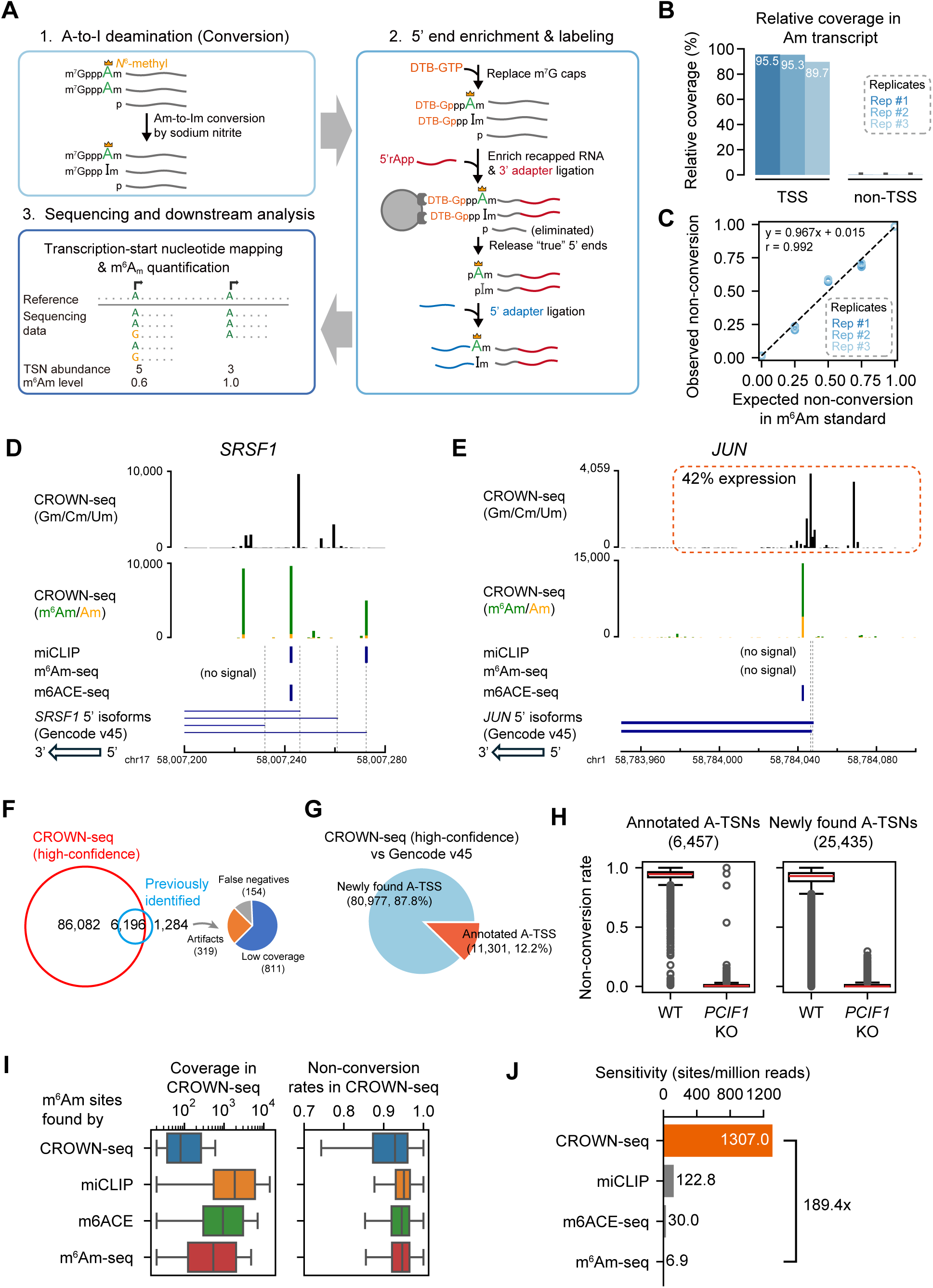
CROWN-seq correctly maps and quantifies m^6^Am. **(A)** Schematic of CROWN-seq. RNAs are firstly treated with sodium nitrite, which causes Am at the transcription-start position to be converted to Im. To isolate the TSN, m^7^G caps are replaced with 3’-desthiobiotin (DTB) caps. These DTB caps are enriched on streptavidin beads, while uncapped background RNA fragments are uncapped and washed away. After washing, an enriched pool of transcript 5’ ends is released from the beads by cleaving the triphosphate bridge, leaving 5’ monophosphate ends that are ligated to an adapter. After adapter ligation, cDNA was synthesized and amplified for Illumina sequencing. During sequencing, the converted sequences were aligned to a reference genome. The TSNs can be determined as the first base immediately after the 5’ adapter sequence. To quantify m^6^Am stoichiometry, we count the number of A (m^6^Am) and G (Am) bases at the TSN position. **(B)** CROWN-seq enriches reads that contain the TSN. The relative coverage of reads mapped to the TSS and non-TSS regions across the m^7^G-ppp-Am-initiated RNA standard was calculated. The average relative coverage of reads that map to the TSS and to non-TSS positions are shown for three replicates. The 95% CI of the relative coverages is shown using error bars. **(C)** CROWN-seq exhibits high quantitative accuracy for measuring m^6^Am stoichiometry. RNA standards (**Table S2**) were prepared with 0%, 25%, 50%, 75%, and 100% m^6^Am stoichiometry. To make m^6^Am standards in different m^6^Am levels, we generated both Am transcripts and m^6^Am transcripts by *in vitro* transcription with cap analogs m^7^G-ppp-Am and m^7^G-ppp-m^6^Am. Five transcripts were made in the Am and m6Am form and mixed to achieve the indicated m^6^Am stoichiometry. These transcripts have identical 5’ ends and different barcodes (**Table S2**). Linear least-squares regression was performed to calculate the correlation between expected non-conversion rates and the observed average non-conversion rates for each standard. All TSNs shown in this plot have high sequencing coverage, ranging from 656 to 21,545 reads. **(D)** CROWN-seq results for *SRSF1*. CROWN-seq shows that 54.0% of *SRSF1* transcripts initiate with A. Among the A-initiated transcripts, 93.4% were resistant to conversion (A’s, shown in green), and therefore m^6^Am. As a result, *SRSF1* has 50.4% m^6^Am transcripts, 3.6% Am transcripts, and 46.0% non-A-initiated transcripts. Notably, a previous miCLIP study identified an internal m^6^A site^44^ which we found was m^6^Am at the TSN based on CROWN-seq. **(E)** CROWN-seq results for *JUN*. CROWN-seq shows that ∼58% of *JUN* transcripts initiate with A. Unlike *SRSF1* which A-TSNs are highly methylated, *JUN* A-TSNs are only ∼75% methylated. As a result, *JUN* has 43.5% m^6^Am transcripts, 14.5% Am transcripts, and 42% non-A-initiated transcripts. **(F)** CROWN-seq identifies most m^6^Am sites identified in previous studies. 7,480 m^6^Am sites in HEK293T cells found either by miCLIP^14^, m^6^Am-seq^11^, or m6ACE-seq^12^ were analyzed. The high-confidence sites in CROWN-seq were defined as A-TSN with ≥20 unique mapped reads. The results shown are from HEK293T cells, which is the same cell line used in all previous studies. Among the 1,284 sites uniquely found in other studies, 811 sites are also mapped by CROWN-seq but at lower coverage (1-19 reads); 319 sites are mapped very far (>100 nt) away from any TSS annotation and thus can be considered as false positives; the remaining 154 sites are mapped very closely to known TSSs and may be false negative results in CROWN-seq. **(G)** Many A-TSNs identified in CROWN-seq in HEK293T cells are not annotated. In this analysis, A-TSSs in **(F)** were intersected with the TSS annotation in Gencode v45. Only 12.2% of A-TSSs found by CROWN-seq are previously annotated. **(H)** CROWN-seq exhibits high accuracy in TSN discovery. In this analysis, we compared the non-conversion of A-TSNs between wild-type and *PCIF1* knockout cells. For the 6,457 A-TSNs annotated by Gencode v45, most of them have high non-conversion rates in wild-type cells and very low non-conversion rates in *PCIF1* knockout cells, indicating correct TSN mapping. Similar to the annotated TSNs, 25,435 newly found A-TSNs were also found to have differential m^6^Am between wild-type and *PCIF1* knockout. Thus, these newly found A-TSNs were also mostly true positives. In this analysis, only A-TSNs mapped by at least 20 reads in both wild-type and *PCIF1* knockout HEK293T cells were used. The details of these sites can be found in **Table S3**. **(I)** The previously identified m^6^Am sites are biasedly in higher expression and higher m^6^Am stoichiometry. Shown are the sequencing coverage (left) and non-conversion rates (right) of different sets of m^6^Am sites in HEK293T CROWN-seq data. In total, 98,147 sites found by CROWN-seq, 2,129 sites found by miCLIP^14^, 3,693 sites found by m6ACE-seq^12^, and 1,610 sites found by m^6^Am-seq^11^ are shown. **(J)** CROWN-seq has much higher sensitivity in m^6^Am discovery than all existing m^6^Am mapping methods. In this analysis, sensitivity is defined as m^6^Am/A-TSN found per million mapped reads. For CROWN-seq, sensitivity was defined as the slope of linear regression result between sequencing depth and A-TSN number among different samples in this study (see **Figure S2G**). For other methods, sensitivity was defined as the number of reported m^6^Am sites over the number of reads in all libraries required for m^6^Am identification.

To increase the accuracy of m^6^Am quantification, we made several optimizations to the ReCappable-seq protocol to markedly increase the read depth of TSNs. These include steps for on-bead adapter ligation and the introduction of unique molecular identifiers (UMIs) in the library preparation (see **Methods**).

### Benchmarking CROWN-seq using m^6^Am-modified standards

To test TSN enrichment in CROWN-seq, we used a m^7^G-ppp-m^6^Am standard spiked into cellular mRNA. Among three replicates, we observed that ∼93% of the reads mapped to the TSN (**Figure 2B**), confirming the enrichment of TSN. To further assess the enrichment of TSNs, we performed GLORI on the same sample. However, in GLORI only a few reads map to the TSN (**Figure S2C**). Thus, the decapping-and-recapping approach markedly enriches for TSNs.

We next wanted to determine the quantitative accuracy of CROWN-seq. To test this, we performed CROWN-seq on a mixture of RNA standards with predefined ratios of m^6^Am and Am (see **Methods** and **Table S2**). We found a highly linear correlation between the expected m^6^Am levels and the observed non-conversion rates measured by CROWN-seq across three replicates (Pearson’s r = 0.992, **Figure 2C**). Taken together, CROWN-seq achieves both precise TSS mapping and m^6^Am quantification in m^6^Am standards.

### CROWN-seq markedly expands the number of mapped m^6^Am sites

To assess the ability of CROWN-seq to map and quantify m^6^Am throughout the transcriptome, we performed CROWN-seq on poly(A)-selected RNA from HEK293T. In total, we identified 219,195 high-confidence TSNs, of which 92,278 were A-TSNs **(Note S2**, “Selection of m^6^Am identification cutoffs”). These TSNs were highly reproducible across biological and technical replicates (**Figure S2E**). Among the A-TSNs, 89,898 were from protein-coding genes, and 219 were from snRNA or snoRNA. Notably, among the mRNA A-TSNs, nearly all had high non-conversion rates (**Figure S2F**), indicating that nearly all A-TSNs contain high stoichiometry m^6^Am.

In contrast to previous m^6^Am mapping methods, CROWN-Seq reveals the diversity of TSNs among all the transcript isoforms for each gene. For example, in the case of *SRSF1*, m^6^Am is readily visible along with multiple other TSNs comprising Gm, Cm, or Um (**Figure 2D**). CROWN-seq also shows that A-TSNs can have intermediate m^6^Am stoichiometry. For example, *JUN* expresses a 5’ transcript isoform with an A-TSN, of which ∼75% of transcript copies are m^6^Am modified (**Figure 2E**). Overall, CROWN-seq provides a comprehensive assessment of all TSNs in a gene and reveals the fraction of each A-TSN that is m^6^Am.

To confirm the accuracy of the mapped m^6^Am TSNs, we examined the 7,480 m^6^Am sites reported by miCLIP^14^, m^6^Am-seq^11^, or m6ACE-seq^12^. Among these sites, the vast majority (∼82.8%, 6,196 of 7,480) were also found among the high-confidence A-TSNs in CROWN-seq (**Figure 2F**). For the remaining 1,284 sites, 811 are also found in CROWN-seq, but in lower sequencing depth; 319 were located far away (>100 nt) from any known TSSs and thus may be artifacts. Thus, CROWN-seq is highly reliable in detecting known m^6^Am sites. The low consistency between previous m^6^Am mapping studies likely reflects incomplete m^6^Am mapping in previous methods.

CROWN-seq clearly identified vastly more A-TSNs than all the other m^6^Am mapping methods combined (12.3-fold, 92,278 vs. 7,480). Notably, only ∼12.2% of the newly found A-TSNs in CROWN-seq are annotated in Gencode v45 (**Figure 2G, S2G**), which primarily relies on CAGE data (see gene annotation guidelines by HAVANA^23^). Notably, the newly identified TSNs with high coverage tend to overlap with or locate proximally to known TSSs, while the ones with low coverage tend to locate further to the known TSSs (**Figure S2G**). The newly identified A-TSNs could be artifacts or could be actual TSNs that were undetected by previous TSS-mapping studies. We suspect that these are true A-TSNs for two reasons: First, as part of the mapping criteria, a minimum of 20 independent reads across all replicates was required for TSN identification. Second, if these sites were RNA fragments, they would not contain m^6^Am. However, these A-TSNs show high stoichiometry of m^6^Am (i.e., non-conversion) in CROWN-seq (**Figure 2H, S2F**) which is lost in *PCIF1* knockout cells (**Figure 2H**), except for some outliers such as TSNs of *S100A6*, *IFI27*, and *ALDH1A1* (see **Table S3**). Thus, the marked increase in the number of m^6^Am sites revealed by CROWN-seq reflects the preferential enrichment for mRNA 5’ ends, which leads to high sensitivity and read depth at TSNs transcriptome-wide.

In contrast to m^6^Am sites identified in CROWN-seq, m^6^Am that were identified in previous m^6^Am mapping methods tended to derive from high abundance transcripts or high abundance TSNs (**Figure 2I**). Because of the high read depth at TSNs, CROWN-seq enables the detection of m^6^Am at more m^6^Am sites than previous methods (**Figure 2J**, **Figure S2H**). Although we used a 20-read cutoff for mapping m^6^Am, m^6^Am sites identified with fewer reads are also likely to represent true TSNs. These m^6^Am TSNs typically show high non-conversion (e.g., 2 or 3 reads among a total of 3 reads) in HEK293T cells but zero non-conversions in *PCIF1* knockout cells (**Figure S2I**). The PCIF1 dependence of these sites is consistent with a true m^6^Am TSN and further highlights the sensitivity of CROWN-seq for mapping m^6^Am at TSNs.

### CROWN-seq reveals consistently high m^6^Am stoichiometry in mRNA across diverse human cell lines

Although our data showed that m^6^Am in mRNA generally exhibits very high stoichiometry (**Figure S2F**), we considered the possibility that these results were unique to HEK293T cells. Several studies have shown that PCIF1 expression can vary considerably in different cell lines^5,24^, which may indicate that m^6^Am stoichiometry is dependent on the cell line. We therefore wanted to determine the m^6^Am landscape across cell lines with varying levels of PCIF1.

We selected several cell lines for this analysis. First, we chose HEK293T, A549, HepG2, and K562 cells, which have also been characterized using multiple orthogonal datasets^25^. Second, we selected colorectal cancer cells (i.e., HT-29 and HCT-116), since PCIF1 depletion in these cells affects their migration and sensitivity to immunotherapy^5^. These colorectal cancer cells have high PCIF1 expression based on western blotting and RT-qPCR, while the non-cancerous colon cell line CCD841 CoN has very low PCIF1 expression^5^. Third, we selected cells with very low CTBP2 expression, a proposed coactivator of PCIF1^24^. These cells, which include Jurkat E6.1 and Huh-7, as well as the previously mentioned K562 and HepG2 cells, would be expected to have low m^6^Am levels based on their low CTBP2 expression^24^ (**Figure S3A**).

For each cell line, we performed CROWN-seq using two to four replicates. In total, we obtained 514 million aligned reads (**Table S3**). In each cell line, 14,650 to 58,768 mRNA A-TSNs with at least 50 reads were analyzed (**Table S3**). The 50-read threshold provides highly consistent quantification of m^6^Am stoichiometry between replicates (**Figure S3B** and **Note S2**).

Quantification of m^6^Am across all TSNs showed that mRNA m^6^Am stoichiometry is generally high. For most of the cells, the average m^6^Am stoichiometry is 0.895±0.03 (**Figure 3A**), indicating high overall mRNA m^6^Am levels. Some cell lines, for example, Jurkat E6.1, HT-29, and Huh-7 cells show very high and less variable m^6^Am levels (0.933±0.1, 0.924±0.1, and 0.916±0.1, respectively); while other cell lines such as CCD841 CoN, HCT-116, and K-562 have relatively low and more variable m^6^Am levels (0.825±0.2, 0.877±0.1, and 0.891±0.1, respectively). It should be noted that in all cell lines, the m^6^Am stoichiometry is still high compared with other mRNA modifications^20^.

**Figure 3.**
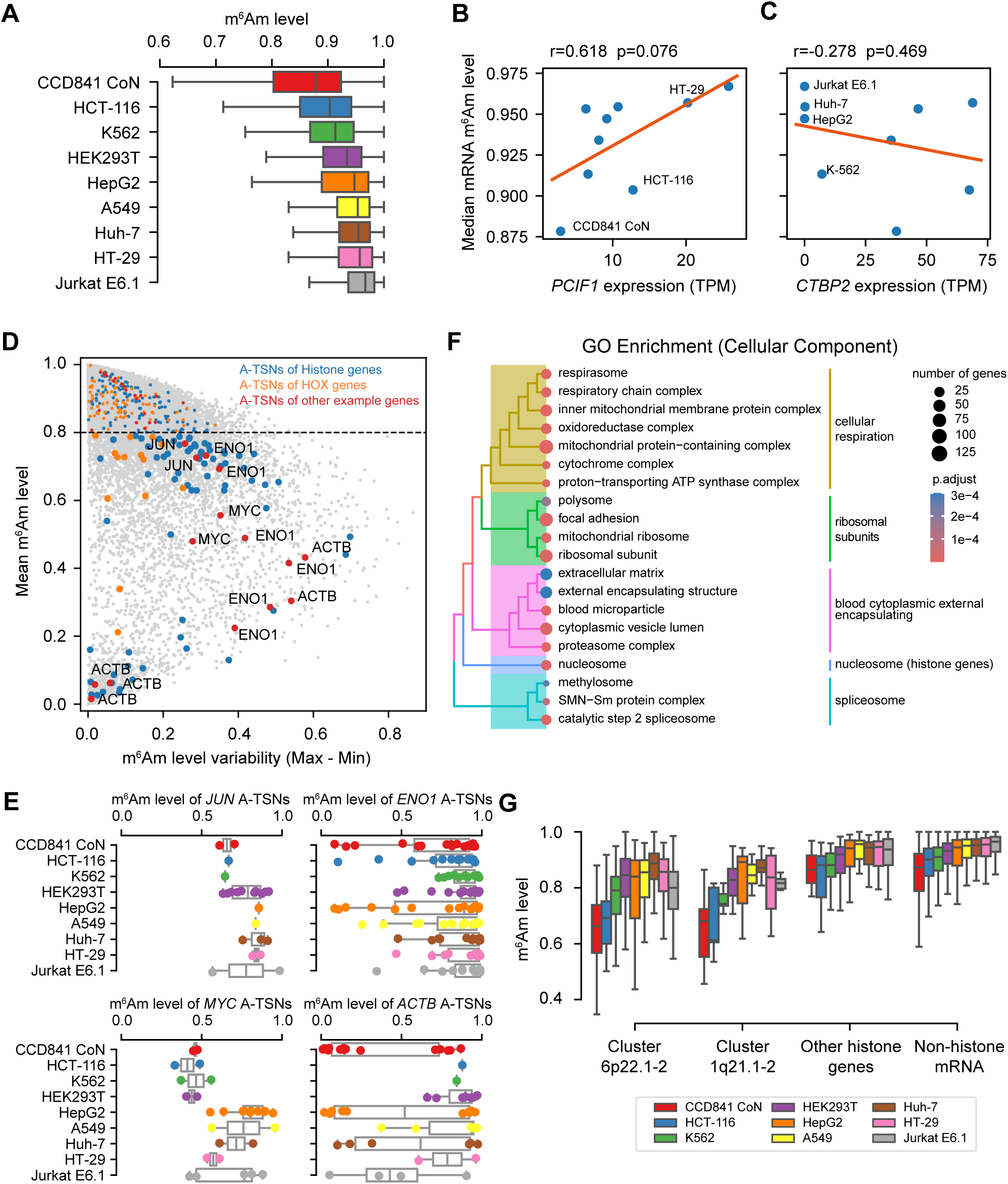
CROWN-seq reveals m^6^Am landscape in mRNA. **(A)** Boxplot showing the overall mRNA m^6^Am levels (i.e., m^6^Am stoichiometry) among different cell lines. The exact m^6^Am levels of the mRNA m^6^Am sites in this analysis can be found in **Table S3**. Only m^6^Am sites with ≥50 reads mapped in at least one cell line were analyzed. **(B)** mRNA m^6^Am stoichiometry is positively correlated with PCIF1 expression. In this plot, *PCIF1* expression was estimated by the number of reads mapped to *PCIF1* TSSs. The read counts were normalized into transcription-start nucleotide per million (TPM) for gene expression comparison. Three cell lines (CCD841 CoN, HCT-116, and HT-29) whose *PCIF1* expression was estimated by western blots and RT-qPCR by Wang *et al.* ^5^ are highlighted. Pearson’s r and p-value in this analysis were obtained by linear regression. **(C)** Overall mRNA m^6^Am stoichiometry is not correlated with CTBP2 expression. Similar to **(B)**, *CTBP2* expression was estimated by CROWN-seq. Four cell lines with very low *CTBP2* expression are highlighted. **(D)** Some A-TSNs have relatively low and more variable m^6^Am stoichiometry among cell lines. In this plot, the variability of the m^6^Am stoichiometry of a site, which is defined as the maximum m^6^Am subtracted by the minimum m^6^Am stoichiometry among all cell lines is shown on the X-axis; the average m^6^Am level of a site among all cell lines is shown on the Y-axis. Several example genes are indicated in different colors. **(E)** Boxplots and dotplots showing the m^6^Am levels of different A-TSNs in *JUN*, *ENO1*, *MYC*, and *ACTB*. These genes contain A-TSNs with relatively low m^6^Am stoichiometry. In this plot, the exact m^6^Am levels of individual A-TSNs are shown in dots, while the median and IQR of the m^6^Am levels are shown in boxplots. Only m^6^Am sites with ≥50 reads mapped were analyzed. **(F)** Gene Ontology enrichment (Cellular Components) of genes containing lowly methylated m^6^Am sites. The full Gene Ontology results can be found in **Table S4**. **(G)** A-TSNs in histone genes tend to have relatively low m^6^Am stoichiometry. In this plot, histone genes are categorized by their genomic localizations. Histone gene cluster 6p22.1-2 and 1q21.1-2 are the two major histone gene clusters. For histone gene cluster 6p22.1-2, 55 to 173 A-TSNs are shown in different cell lines; for histone gene cluster 1q21.1-2, 9 to 14 A-TSNs are shown; for other histone genes, 24 to 109 A-TSNs are shown.

We considered the possibility that the high m^6^Am stoichiometry might be caused by RNA structure that blocks access to sodium nitrite leading to non-conversion. However, essentially complete conversion was seen in *PCIF1* knockout HEK293T cells, which makes it likely that m^6^Am is the cause of non-conversions. Also, we found that A-TSNs completely converted in 5’ ends predicted to be highly structured, suggesting that RNA structure does not impair conversion in CROWN-seq (**Figure S3C**).

The differences in m^6^Am stoichiometry are related to PCIF1 expression (**Figure 3B, S3D**). For example, CCD841 CoN cells, which have very low PCIF1 expression based on our measurements (**Figure 3B, S3D**) and previous measurements^5^, exhibit the lowest median m^6^Am stoichiometry at ∼0.878. However, even this stoichiometry is still relatively high. Thus, m^6^Am levels are affected by PCIF1 expression, but m^6^Am can be considered as a high stoichiometry modification across all tested cell lines. On the other hand, the proposed PCIF1 coactivator CTBP2, exhibited a weak correlation to mRNA m^6^Am (**Figure 3C**).

### Several mRNAs show low m^6^Am stoichiometry

Although most A-TSNs in mRNA exhibit high m^6^Am stoichiometry, some exhibit stoichiometry below 0.8, and even below 0.5 (**Figure 3A** and **Table S3**). To identify A-TSNs with low m^6^Am, we examined each A-TSN and calculated its average stoichiometry across all cell lines (**Figure 3D**). For each A-TSN we also assessed its variability by calculating the range of m^6^Am levels measured across cell lines (**Figure 3D**). This analysis demonstrates that a significant subset of A-TSNs have low stoichiometry, with some showing variability depending on the cell type. For example, *JUN* contains a lowly methylated A-TSN, as shown above in HEK293T cells (**Figure 2E**), and also exhibits low stoichiometry in many other cell lines (**Figure 3E**). In addition, *ENO1*, *MYC*, and *ACTB* also show low m^6^Am stoichiometry in some of their A-TSNs (**Figure 3E**).

We next used Gene Ontology (GO) analysis to determine if the low m^6^Am A-TSNs are associated with specific cellular functions. The GO analysis of Cellular Component categories showed a marked enrichment of genes linked to cellular respiration, ribosomal subunits, spliceosome, and nucleosome (which are mostly histone genes) (**Figure 3F**, **Table S4**). Similar results were found in the Biological Processes GO analysis (**Table S4**). In addition to these genes, we also noticed HOX genes contain lowly methylated A-TSNs (**Figure S3E**).

Among all different gene categories, histone genes exhibited the lowest overall m^6^Am stoichiometry (**Figure 3G**). Notably, histone genes have unique mechanisms of gene expression. Many histone genes are located in gene clusters (i.e., clusters 6p22.1-2 and 1q21.1-2) and transcribed in histone locus bodies^26^. These clustered histone genes tend to contain upstream TATA-box and downstream T-rich sequences (**Figure S3F**). In contrast, non-clustered histone genes tend to have high m^6^Am stoichiometry (**Figure 3G**) and show different promoter sequence contexts (**Figure S3F**). This data suggests that transcription mechanisms might be important for determining m^6^Am stoichiometry.

### m^6^Am stoichiometry is linked to the sequence of core promoter

The differential methylation in histone genes based on their transcription mechanisms raises the possibility that transcription initiation mechanisms might affect m^6^Am stoichiometry. Because m^6^Am is the first nucleotide in mRNA, its deposition may be highly influenced by early transcription events. Notably, PCIF1 binds to RNA polymerase II^27^ and is enriched in promoter regions^28^, which may be important for methylation of the 5’ end of mRNAs. We therefore considered the possibility that different transcription mechanisms may be linked to different levels of m^6^Am.

As a first test, we examined whether nucleotide preferences upstream (which would reflect sequences involved in transcription initiation) or downstream of the A-TSN are linked to m^6^Am stoichiometry. We binned A-TSNs based on the m^6^Am stoichiometry and examined nucleotide preferences at each position. Using this approach, we found that the nucleotides upstream of the A-TSN were markedly different for A-TSNs with low vs. high m^6^Am stoichiometry (**Figure S4A**). For example, at positions -4 and -1, there was a clear positive correlation between the use of C and m^6^Am stoichiometry (**Figure 4A**). The correlation of these nucleotide positions that lie in the promoter region to m^6^Am stoichiometry suggests that transcriptional mechanisms might influence m^6^Am stoichiometry.

**Figure 4.**
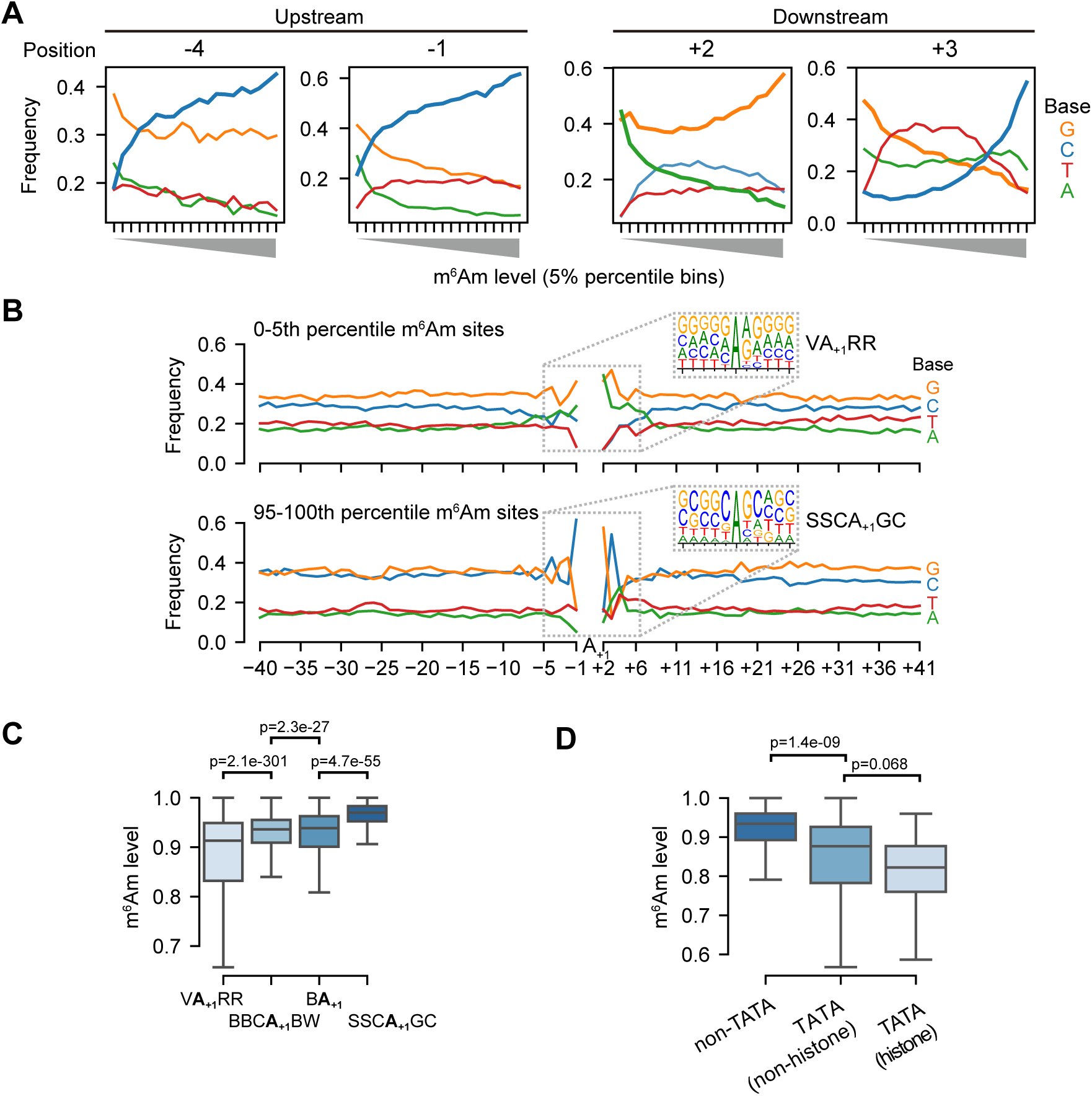
m^6^Am stoichiometry is related to core promoter sequence. **(A)** m^6^Am stoichiometry is related to base composition in both upstream and downstream of A- TSNs. In these plots, 58,723 A-TSNs are grouped into twenty 5-percentile bins (X-axis). For each bin, the frequency of A, T, C, and G bases at each position relative to the A-TSN are plotted on the Y-axis. Among different positions in the promoter region, C’s in -4, -1, and +3, as well as G’s in +2 are positively correlated with high m^6^Am; while A’s in +2 are negatively correlated with high m^6^Am. Results for other promoter positions can be found in **Figure S4A**. **(B)** Motif analysis of A-TSNs with the lowest 5% m^6^Am stoichiometry (upper) and the A-TSNs with the highest 5% m^6^Am stoichiometry (lower). The core promoter region (-40 to +41) was screened for enriched motifs. The lowest 5% A-TSNs exhibited a VA_+1_RR TSS (V=A/C/G, R=A/G) motif, while the highest 5% A-TSNs exhibited a SSCA_+1_GC (S=C/G) motif. The sequence contexts for all A-TSNs are shown in **Figure S3E**. **(C)** A-TSNs expressed from different core promoters exhibit different m^6^Am stoichiometry. Core promoters containing the VA_+1_RR motif produce transcripts with relatively low m^6^Am stoichiometry. Transcripts using the SSCA_+1_GC motif exhibited relatively high m^6^Am stoichiometry. In comparison, the m^6^Am stoichiometry in conventional A-TSNs from either BBCA_+1_BW or BA_+1_ is also shown and exhibits intermediate m6Am stoichiometry. In this analysis, 14,788, 7,981, 34,578, and 1,376 A-TSNs were used for each of the four motifs. P-values, Student’s t-test, two-sided. **(D)** TATA-box containing core promoters exhibit relatively low m^6^Am stoichiometry. For this analysis, the TATA-box is defined as TATAWAWR^29^. Because many TATA-boxes found in our A-TSN dataset are outside the classic -31 to -24 region, we extended the region for the TATA-box search to -36 to -19. Since histone genes preferentially contain TATA box, we separately plotted TATA-box-containing histone genes (N=155) and TATA-box-containing non-histone genes (N=28). 58,540 A-TSNs without TATA-box are also shown. P-values, Student’s t-test, two-sided.

We also observed strong nucleotide preferences at positions downstream of the A-TSN. These include nucleotide preferences at +2 (**Figure 4A**). These could reflect sequence preferences for PCIF1; however, this position is also part of transcription-initiation motifs (see below), and thus the contribution of transcription mechanisms and direct sequence preferences of PCIF1 are difficult to deconvolve.

To more directly determine specific transcription mechanisms linked to m^6^Am, we examined how specific sequence motifs around A-TSNs correlate with m^6^Am stoichiometry. We found markedly different sequence motifs surrounding highly methylated A-TSNs compared to lowly methylated A-TSNs (**Figure 4B**). A-TSNs with the highest m^6^Am stoichiometry (top 5th-percentile, 0.991 average stoichiometry) are enriched in an SSCA_+1_GC (S=C/G) motif, which is similar but distinct from the well-known BBCA_+1_BW (B=C/G/T, W=A/T) transcription initiator motif^29^, largely because of the C at the +3 position (**Figure 4B**). In contrast, the A-TSNs with the lowest m^6^Am stoichiometry (bottom 5th-percentile, 0.435 average stoichiometry) were enriched in an unconventional VA_+1_RR (V=A/C/G, R=A/G) motif (**Figure 4B**).

We next classified each A-TSN based on whether they use the SSCA_+1_GC or VA_+1_RR motifs, or if they contain the conventional BBCA_+1_BW and BA_+1_ Inr-like motifs (**Figure S4B**). This plot shows that BBCA_+1_BW and BA_+1_ motifs exhibit intermediate m^6^Am stoichiometry (**Figure 4C**). Overall, these data indicate that m^6^Am stoichiometry is strongly related to the TSS motif in the core promoter, which implies that m^6^Am formation is linked to the transcription initiation process.

Because transcription initiation is also affected by other elements in the core promoter^29^, we also asked whether these transcription-related elements, such as TATA-box and transcription factor-binding sites, are associated with higher or lower m^6^Am stoichiometry. We first analyzed the relationship between m^6^Am and elements including the TATA-box, BREu, BREd, and DCE^29^. In this analysis, A-TSNs from promoters containing TATA-box exhibited lower m^6^Am stoichiometry, especially those of histone genes (**Figure 4D**). On the other hand, other elements, such as BREu and BREd, which are motifs for recruitment of TFIIB^29^, and DCE, which binds by TAF1^29^, showed little correlation with m^6^Am stoichiometry (**Figure S4C**). Thus, the presence of the TATA box exhibited the strongest effect and predicted lower m^6^Am stoichiometry.

We next analyzed the relationship between m^6^Am and transcription factor-binding sites (TFBS). To test this, we screened A-TSNs for the presence of 401 transcription-factor binding sites and examined the relationship between the binding sites and m^6^Am stoichiometry (see **Methods**). Several TFBSs, such as those for NANOG and FOXJ3, exhibited a slight negative correlation to m^6^Am (**Figure S4D**); while other TFBS, such as SP2 and KLF4, exhibited a slight positive correlation to m^6^Am (**Figure S4E**). Overall, no specific TFBS exhibited a strong effect on m^6^Am stoichiometry (**Figure S4F**).

Taken together, our data show a linkage between transcriptional mechanisms and m^6^Am stoichiometry.

### m^6^Am does not substantially influence mRNA stability or translation

Previous studies sought to determine the effect of m^6^Am on mRNA stability and translation based on gene-level annotations of the starting nucleotide^4,14,15,30^. However, the gene level annotations do not take into account the potential for many transcription-start nucleotides (**Figure S5**). Rather than using a binary metric of m^6^Am or non-m^6^Am, we developed a metric that reports the fraction of all TSNs for each gene that contain m^6^Am. This “m^6^Am gene index” is the ratio of m^6^Am TSNs over all TSNs, as measured by CROWN-seq, for each gene. Using the m^6^Am gene index, we reanalyzed the previously published translation efficiency^4,14^ and RNA stability^14^ data in HEK293T cells. We found that genes with low or high m^6^Am gene index do not show differences in translation (**Figure S6A, S6B**) or RNA stability (**Figure S6C**) in *PCIF1* knockout cells compared to wild-type.

### m^6^Am is involved in efficient transcription of A-initiated transcripts

We next wanted to examine other potential functions of m^6^Am. Although we found no clear effect of m^6^Am on mRNA stability, we asked if m^6^Am influences transcript expression levels. To test this, we quantified the abundance of each A-TSN isoform in HEK293T and A549 cells. For these experiments, we added a mixture of pre-capped ERCC spike-ins (see **Methods**) to the RNA samples before performing TSN expression quantification by ReCappable-seq. This ERCC spike-in mixture calibrates sequencing results and increases TSN expression quantification accuracy (see **Methods**).

In this analysis, we binned A-TSNs into percentiles based on their m^6^Am stoichiometry. Here we could see that transcripts with the highest levels of m^6^Am also exhibited the highest overall expression levels (**Figure 5A**, **S6D**, and **Table S5**). This suggests that m^6^Am is associated with higher transcript expression.

**Figure 5.**
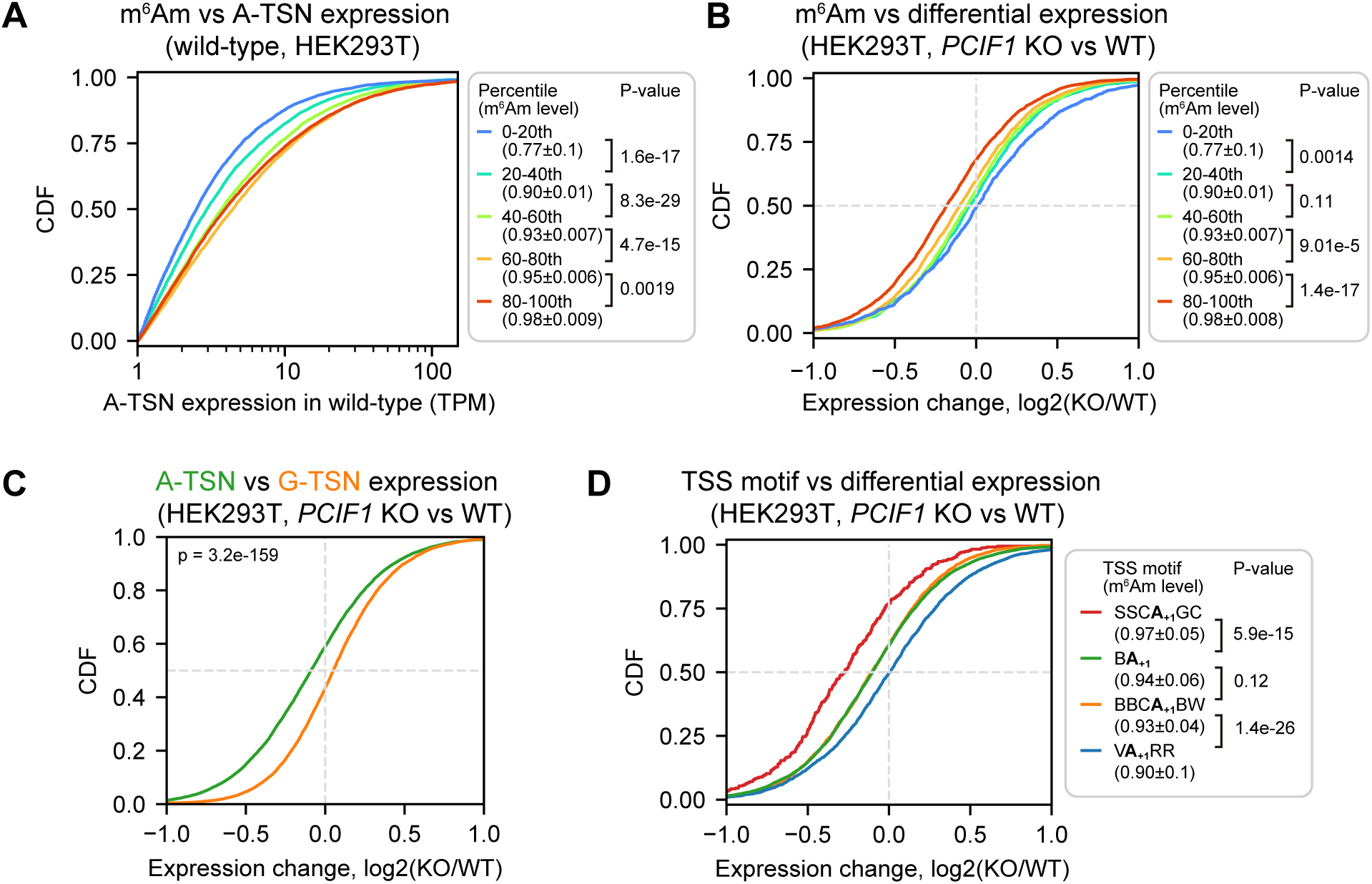
*PCIF1* knockout leads to m^6^Am and TSS motif-related A-TSN expression changes. **(A)** m^6^Am stoichiometry is positively related to A-TSN expression in wild-type HEK293T cells. In this cumulative distribution plot, the expression of each A-TSN was quantified by ReCappable-seq, for all A-TSNs in each indicated m^6^Am stoichiometry bin. A-TSNs (n=58,723) were grouped into five bins based on m^6^Am stoichiometry quantified by CROWN-seq. In total, 5,125, 6,962, 7,991, 8,368, and 8,009 A-TSNs are shown in each bin (from low m^6^Am to high m^6^Am). These A-TSNs have an average TPM ≥1 in two ReCappable-seq replicates and coverage ≥50 in CROWN-seq. P-values, Student’s t-test for TPM (log-transformed), two-sided. **(B)** The expression level of high m^6^Am stoichiometry A-TSNs is reduced in *PCIF1* knockout. Shown is a cumulative distribution plot of A-TSN expression change in HEK293T cells upon *PCIF1* knockout. The differential expression of A-TSN was calculated by DESeq2^45^. Similar to **(A)**, the A-TSNs were binned based on the m^6^Am stoichiometry. In total, 3,269, 2,272, 3,218, 3,813, and 3,369 A-TSNs are shown in each bin (from low m^6^Am to high m^6^Am). A-TSNs with a baseMean (i.e., the average of the normalized count among replicates) ≥100 were used in the differential expression test (two replicates were used for both wild-type and *PCIF1* knockout) and coverage ≥50 reads in CROWN-seq. P-values, Student’s t-test, two-sided. **(C)** Shown are cumulative distribution plots of expression changes of A-TSNs and G-TSNs after PCIF1 depletion. 14,516 A-TSNs and 9,667 G-TSNs with expression levels quantified by ReCappable-seq are shown. These A-TSNs and G-TSNs have baseMean ≥100 during the differential expression test. P-values, Student’s t-test, two-sided. **(D)** Similar to **(B)**, A-TSNs that use different TSS motifs exhibit different changes in expression upon *PCIF1* knockout. In total, 352 A-TSNs using SSCA_+1_GC, 7,928 A-TSNs using BA_+1_, 2,958 A-TSNs using BBCA_+1_BW, and 2,760 A-TSNs using VA_+1_RR are shown. These A-TSNs have baseMean≥100 during differential expression test (2 replicates for both wild-type and *PCIF1* knockout) and coverage ≥50 reads in CROWN-seq. P-values, Student’s t-test, two-sided.

To determine if m^6^Am causes the increased expression of A-TSN transcripts, we measured the expression change for each A-TSN in wild-type and *PCIF1* knockout HEK293T and A549 cells (**Table S5**). We found that A-TSNs with higher m^6^Am stoichiometry exhibit significantly reduced expression in *PCIF1* knockout, while A-TSNs with the lowest m^6^Am stoichiometry were almost unchanged (**Figure 5B, S6E**). In contrast, G-TSNs were slightly increased in *PCIF1* knockout cells (**Figure 5C, S6F**). These data suggest that m^6^Am promotes the expression of A-TSN transcripts.

We were surprised that PCIF1 depletion leads to a selective decrease in the expression of A- TSN transcripts in the highest percentile bin but had little to no effect in the other bins. Each bin has very high m^6^Am stoichiometry (∼0.77 in the lowest bin and ∼0.98 in the highest bin in HEK293T) (**Figure 5B, S6E**). Thus, if m^6^Am is a stabilizing mark, we should see reduced expression in all bins. We therefore considered other possibilities that might explain why PCIF1 depletion affects transcript levels in some bins but not others.

An important difference between A-TSN in different bins is that they tend to use different TSS motifs (see **Figure 4C**). We therefore asked if the effect of m^6^Am depletion is linked to the TSS motifs. For this analysis, we classified A-TSNs based on the presence of SSCA_+1_GC, VA_+1_RR, or other TSS motifs (i.e., BBCA_+1_GC and BA_+1_). Here we found that the identity of the TSS motif was highly associated with the degree of transcript reduction in *PCIF1* knockout cells (**Figure 5D, S6G**). Notably, transcripts that use the SSCA_+1_GC motif showed the largest drop in expression. In contrast, A-TSNs that use the VA_+1_RR TSS motif showed almost no change in expression in *PCIF1* knockout cells (**Figure 5D, S6G**).

Taken together, these data suggest that the effect of m^6^Am is not related to mRNA stability but instead is related to transcription. Our data suggest that certain transcription initiation complexes, such as those that use the SSCA_+1_GC motif, rely on m^6^Am for efficient expression. However, other TSS motifs do not rely as strongly on m^6^Am to achieve efficient expression. These data suggest that m^6^Am may have important roles in the transcription processes.

### CROWN-seq reveals diverse m^6^Am stoichiometry in snRNA and snoRNA

In addition to mRNAs, m^6^Am is also found on snRNAs and snoRNA^10,12^. However, the stoichiometry and dynamics of m^6^Am in these RNAs are unknown. Using CROWN-seq we quantified m^6^Am stoichiometry in several snRNAs, including U1, U2, U4, U4ATAC, U5, U7, U11, and U12. These snRNAs are transcribed by RNA polymerase II^31^, are capped, and use A-TSNs^10^. Among these snRNAs, we identified 51 m^6^Am sites, of which 29 were unannotated 5’ variants located close to the annotated TSNs (**Table S6**).

Compared with mRNA, m^6^Am in snRNA exhibited a very different distribution of stoichiometry (**Figure S2F**). First, snRNA m^6^Am sites exhibited generally low m^6^Am stoichiometry, typically below 0.3. Second, m^6^Am stoichiometry between different snRNA genes was much more variable than in mRNA (**Figure 6A**). Third, some snRNA genes show highly variable stoichiometry in different cell lines.

**Figure 6.**
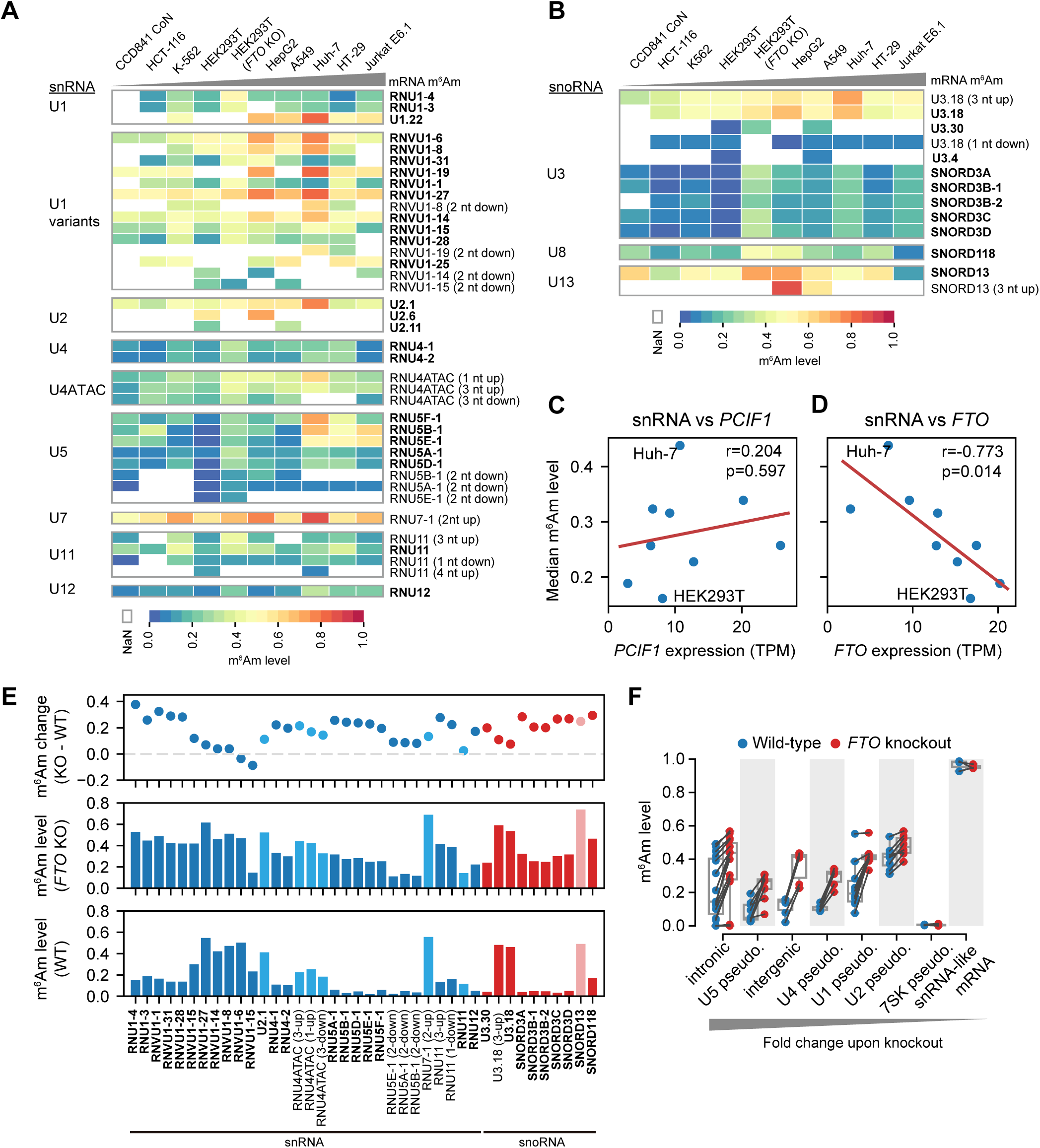
CROWN-seq reveals m^6^Am landscape in snRNA and snoRNA. **(A)** Heatmaps showing m^6^Am stoichiometry in different snRNA gene families and isoforms. Cell lines in the column are ranked by the overall mRNA m^6^Am stoichiometry. The name of each snRNA isoform is shown on the right. A-TSNs already annotated in Gencode v45 are highlighted in bold. For newly found A-TSNs, the relative distance between the new A-TSN and the nearest annotated A-TSN is showed in brackets. Note that Gencode v45 contains snRNA annotation from different databases. For example, *RNU1-4* and *U1.22* are both U1 snRNA, however, *RNU1-4* is from the HGNC database and *U1.22* is from the RFAM database. **(B)** Similar to **(A)**, Heatmaps show the m^6^Am stoichiometry in different snoRNA isoforms. **(C)** and **(D)**, snRNA methylation levels are not well correlated with *PCIF1* expression, but negatively correlated with *FTO* expression. The RNA expression levels of *PCIF1* and *FTO* were estimated by reading counts in CROWN-seq, which were converted into TPM to normalize the sequencing depth. Linear regressions were performed to obtain Pearson’s r and p-value of the correlations. **(E)** FTO depletion leads to increased m^6^Am level (i.e., m^6^Am stoichiometry) in many kinds of snRNA and snoRNA. In this plot, the difference in m^6^Am levels between wild-type and *FTO* knockout cells is shown in the first row. The exact m^6^Am levels in *FTO* knockout and wild-type cells are shown in the second and third rows. Different kinds of snRNA and snoRNA are shown in different colors. **(F)** FTO depletion leads to increased m^6^Am stoichiometry in snRNA and snoRNA pseudogenes. In this plot, shown are the annotated pseudogenes of U1, U2, U4, U5, and 7SK, as well as the newly identified snRNA/snoRNA pseudogenes in intronic and intergenic regions. Several mRNAs exhibited 5’ ends resembling snRNA pseudogenes. However, these snRNA-like mRNA 5’ ends showed high and stable m^6^Am stoichiometry in both wild-type and *FTO* knockout cells.

For example, among U1 snRNA genes, *U1.22* exhibited relatively high m^6^Am levels (∼0.47-0.80) in all cell lines, while *RNU1-3* and *RNU1-4* show relatively low m^6^Am levels (∼0.09-0.45, **Figure 6A** and **Table S6**). U5 snRNAs have the highest variability m^6^Am stoichiometry between cell types (**Figure 6A**). The U5 snRNA isoform *RNU5E-1* varies 31.6-fold in the m^6^Am level between HEK293T (0.0185) and Jurkat E6.1 cells (0.586). In contrast, m^6^Am in U2 and U7 snRNA are generally high (at 0.35-0.76 and 0.48-0.89, respectively) and not variable between cell lines (**Figure 6A**).

In addition to snRNA, we also examined 17 A-TSNs in C/D box snoRNA including U3, U8, and U13. These snoRNAs are involved in rRNA 2’-*O*-methylation during pre-rRNA processing^32^. m^6^Am stoichiometry in snoRNA is highly related to snoRNA species and snoRNA isoform. For example, among different U3 snoRNA isoforms, A-TSNs of *U3.18* have much higher m^6^Am stoichiometry than others (e.g., *SNORD3A*) (**Figure 6B**). These data indicate that snoRNA also has highly variable m^6^Am stoichiometry.

### FTO specifically controls m^6^Am stoichiometry in snRNA and snoRNA

We next sought to understand why m^6^Am stoichiometry is low in snRNA and snoRNA. We considered the possibility that the overall m^6^Am stoichiometry in snRNA is correlated with PCIF1 expression, as we saw with mRNA (**Figure 3B**). However, we found a poor correlation of overall m^6^Am stoichiometry in snRNA with PCIF1 expression (Pearson’s r = 0.204, P-value=0.597, **Figure 6C**).

We next considered FTO, a highly efficient demethylase for m^6^Am in snRNA^10,12^. In contrast to PCIF1, FTO expression exhibited a strong negative correlation with snRNA methylation levels (Pearson’s r = -0.773, P-value=0.014, **Figure 6D**). Notably, HEK293T cells, which were tested in our previous study^10^, exhibited the highest FTO expression and the lowest snRNA m^6^Am stoichiometry (**Figure 6D**). Some other cell lines, such as Huh-7, have lower FTO expression and thus have relatively higher m^6^Am stoichiometry in snRNAs (**Figure 6D**).

We next wanted to determine how FTO affects m^6^Am stoichiometry in snRNAs. Using CROWN-seq on *FTO* knockout HEK293T cells, we observed prominent m^6^Am level increases in nearly all snRNA and snoRNA (**Figure 6E**). Most of the snRNA isoforms exhibited an overall increase in m^6^Am stoichiometry by ∼0.2 upon *FTO* knockout. However, a notable subset of snRNAs were not affected by FTO depletion. For example, the *RNVU1-8* isoform has little change in m^6^Am stoichiometry. *RNVU1-8* has an unusually high m^6^Am stoichiometry at ∼0.47 compared to other U1 snRNA isoforms in wild-type cells (**Figure 6E**).

Notably, FTO depletion does not increase m^6^Am levels in snRNA and snoRNA to the levels seen in mRNA (i.e. >0.9 stoichiometry). This suggests that the low m^6^Am levels in snRNA and snoRNA are not solely due to FTO-mediated demethylation. Instead, these snRNAs are likely to be inefficiently methylated by PCIF1 and are then demethylated by FTO in order to achieve their overall low m^6^Am stoichiometry.

We also found FTO demethylates m^6^Am in snRNA pseudogenes. Overall, we mapped 69 A- TSNs in annotated snRNA/snoRNA pseudogenes. These pseudogenes exhibited increased methylation upon *FTO* knockout (**Figure 6F** and **Table S6**). We also identified 202 snRNA/snoRNA pseudogene-like transcripts (see **Table S6**). These transcripts exhibited very high similarity to the annotated snRNA/snoRNA pseudogenes, and therefore likely reflect previously unannotated pseudogenes (see **Methods**). Upon *FTO* knockout, A-TSNs in these unannotated pseudogenes also exhibited increased m^6^Am levels (**Figure 6F**).

### FTO has minimal effects on m^6^Am and m^6^A at 5’ ends of mRNA

We next asked whether FTO levels affect m^6^Am levels in mRNA. To address this question, we compared FTO RNA expression and median mRNA m^6^Am stoichiometry in all nine cell lines. This analysis shows a weak negative correlation between FTO expression and mRNA m^6^Am (Pearson’s r = -0.239, P-value=0.535, **Figure S7A**).

To further assess whether FTO affects m^6^Am levels in mRNA, we quantified m^6^Am level changes in mRNA in wild-type and *FTO* knockout HEK293T cells. Overall, we observed a very small increase in mRNA m^6^Am with only a few m^6^Am sites having notably increased methylation level upon *FTO* knockout (**Figure S7B**). Thus, only select m^6^Am sites in mRNA are efficiently demethylated by FTO.

Although CROWN-seq focuses on m^6^Am measurements, the reads in CROWN-seq can contain internal m^6^A sites if they are close to the TSN. m^6^A sites are readily detected because they do not undergo conversion with sodium nitrite. We therefore examined the stoichiometry of these 5’-proximal m^6^A sites in *FTO* knockout HEK293T cells. We identified internal m^6^A sites that were mapped with at least 50 reads in both wild-type and *FTO* knockout cells and had a non-conversion rate of ≥0.2 in either genotype. In total, we identified 235 high-confidence m^6^A sites which were found by both CROWN-seq and GLORI^20^(**Table S6**). These m^6^A sites exhibited the expected DRm^6^ACU motif (**Figure S7C**). However, these sites only showed small changes in non-conversion rates (P-value=0.00037, paired t-test) (**Figure S7D** and **Figure S7E**). It should be noted that our conclusion about the effect of FTO on internal m^6^A is restricted to specific m^6^A sites around 5’ ends since most internal m^6^A sites are not found in the 5’ fragments examined in CROWN-seq.

Taken together, FTO has a strong preference for demethylating m^6^Am in snRNA, snoRNA, and their pseudogenes, compared to mRNA. FTO is a major determinant of the overall m^6^Am levels of these transcripts in different cell lines.

## DISCUSSION

Functional studies of m^6^Am require highly accurate transcriptome-wide maps. However, m^6^Am mapping studies have relied on the assumption that each gene can be considered to have a single start nucleotide. To overcome this, we developed CROWN-seq, which maps the TSNs for all 5’ transcript isoforms, and measures the exact stoichiometry of m^6^Am at all A-TSNs.

CROWN-seq reveals a markedly distinct distribution of m^6^Am than previously recognized, largely due to inaccuracies in previous maps, and the problem with assigning each gene to a single start nucleotide. In addition, the quantitative measurements of m^6^Am in CROWN-seq show that the earlier idea that many mRNAs contain transcription-start nucleotide Am is largely incorrect since nearly all A-TSNs exhibit high stoichiometry m^6^Am. Overall, this study establishes the first quantitative, transcript isoform-specific m^6^Am map in mammalian cells. The m^6^Am maps reveal that m^6^Am is associated with increased transcript abundance, with functions of m^6^Am more correlated with transcription initiation than stability.

By selectively capturing and examining only 5’ ends, CROWN-seq achieves exceptional read depth at the TSN, enabling highly accurate identification and quantification of m^6^Am. Notably, CROWN-seq is an antibody-free method and thus avoids the problem of immunoprecipitation of both m^6^Am- and m^6^A-containing fragments. This dual-specificity of antibodies creates ambiguities in m^6^Am mapping. Additionally, antibody binding cannot provide quantitative measurements of m^6^Am. In contrast, CROWN-seq uses a sodium nitrite-based chemical method for m^6^Am identification, which we show fully converts Am to Im, but leaves m^6^Am intact. Thus, the fraction of A-TSNs that contain m^6^Am or Am can be readily determined by sequencing, where all Am nucleotides are read as G. The exceptional read depth of CROWN-seq enables quantification of m^6^Am at single-nucleotide resolution, resulting in vastly more m^6^Am sites than all previous m^6^Am mapping methods combined. Although CROWN-seq involves many chemical and enzymatic steps, m^6^Am quantification by CROWN-seq is very accurate and robust, which was examined by m^6^Am standards, consistency across different replicates, and *PCIF1* knockout data. Notably, chemical conversion-based methods tend to have artifacts in regions with stable RNA secondary structures^33,34^, where the nucleotides cannot efficiently interact with the chemical reagent. However, we found that CROWN-seq is very reliable even for highly structured 5’ ends (**Figure S3C**), which might be due to the high accessibility of the TSN, the high stringency of the conversion steps^20^, or RNA denaturation due to glyoxal^35^.

It is worth mentioning that there is no golden standard for transcription-start nucleotide (site) mapping accuracy estimation. For CROWN-seq, we first tested the mapping accuracy by *in vitro* transcribed RNA oligos, which shows that ∼93% of the 5’ ends can be mapped correctly. However, in practice, *in vitro* transcription might initiate at non-specific TSSs, resulting in 5’ ends not overlapping with the desired TSSs^36,37^. Thus, the mapping accuracy can be underestimated in this assay. Since mRNA A-TSNs in the cells are known to be highly methylated by PCIF1^1,4^, we considered that the presence or absence of m^6^Am at mapped A-TSNs can be used to assess the accuracy of TSN identification. True A-TSNs should have m^6^Am. In CROWN-seq essentially all previously annotated A-TSNs and newly found A-TSNs exhibited high non-conversion rates. These A-TSNs were well converted upon *PCIF1* knockout. This indicates very high TSN mapping accuracy, even at the many previously unannotated TSNs described here. These previously unannotated TSNs were likely missed because traditional transcription-start mapping methods and pipelines lack the sensitivity to discover them. These unannotated TSNs might have specific molecular functions. Future studies might focus on the biology of these unannotated TSNs, for example, whether these unannotated TSNs, compared to major TSNs, are associated with different mRNA processing events, such as alternative splicing.

We performed CROWN-seq in nine different cell types with the goal of understanding common principles that guide m^6^Am formation in mRNA. In all cell types, m^6^Am was a very high stoichiometry modification, with some exceptions. We found a correlation between PCIF1 expression and m^6^Am stoichiometry, but even cells with very low PCIF1 expression exhibited high m^6^Am stoichiometry. The CROWN-seq data is highly consistent with recent mass spectrometry analysis of mRNA caps by us^1^ and others^2,4^. These mass spectrometry studies purified the entire cap structure comprising the m^7^G, the triphosphate linker, and the first nucleotide. In these studies, m^7^G-ppp-m^6^Am was highly prevalent while m^7^G-ppp-Am abundance was typically 1/10 as m^7^G-ppp-m^6^Am in nearly all cell lines^1,2^. This mass spectrometry data was the first suggestion that transcription-start nucleotide Am was not a prevalent modification in mRNA, as had been suggested by early chromatography studies^9^. We suspect that the high levels of Am seen in these early analyses of mRNA can be explained by contaminating snRNA or rRNA fragments, which are highly difficult to remove, even with multiple rounds of poly(A) purification^38^. It remains possible that there are cell types or cellular contexts that remain to be discovered with low m^6^Am (i.e., high Am) levels. However, it is clear that high m^6^Am stoichiometry is a general feature of most or all cell types in this study.

The initial m^6^Am maps relied on published TSN annotations. In the first m^6^Am map, annotations were based on FANTOM5^16^, which primarily uses CAGE datasets to define the start nucleotide. However, these annotations selected a single start nucleotide even if multiple TSS signals from CAGE peaks were detected for a gene^4,14^. It should be noted that some genes may primarily use m^6^Am for all 5’ transcript isoforms. These genes would therefore have a high m^6^Am gene index. Genes with a high m^6^Am gene index are likely to be preferentially affected by PCIF1 depletion or pathways that affect m^6^Am.

Based on the small range of m^6^Am stoichiometry in A-initiated mRNAs, it is unlikely that the variability in stoichiometry has functional significance for most mRNAs. Instead, our data suggest that mRNAs initiate with either Gm, Cm, Um, or Am, where Am is highly m^6^Am modified. mRNAs that initiate with m^6^Am may have shared regulatory mechanisms that distinguish them from mRNAs that initiate with Gm, Cm, and Um. Additionally, genes that primarily initiate with m^6^Am, either because they have only one major transcription-start site, or because all their transcription-start nucleotides are A, would be highly influenced by m^6^Am-dependent regulatory mechanisms. Currently, cellular pathways that target m^6^Am-initiated mRNAs are not well known.

Our study revealed a link between m^6^Am and transcription. This effect was detectable because of the highly quantitative nature of m^6^Am measurement in CROWN-seq. Although all A-TSNs show high stoichiometry, there are differences in the overall m^6^Am stoichiometry between transcripts, e.g., ∼0.85 stoichiometry vs. 0.95 stoichiometry which can readily be detected by CROWN-seq. We found that these differences are often related to the specific TSS motif. For example, the Inr-like SSCA_+1_GC TSS motif was associated with the highest m^6^Am stoichiometry, while transcripts using the VA_+1_RR TSS motif exhibited relatively lower m^6^Am stoichiometry. This finding highlights that the major role of m^6^Am might be linked to transcription regulation, which is supported by a recent study by An et al.^39^.

We then examined the effects of PCIF1 depletion on m^6^Am transcript abundance. We found that transcripts with higher methylation in wild-type cells tend to have a larger reduction in RNA expression level upon *PCIF1* knockout. Further analysis showed that transcripts that use the SSCA_+1_GC TSS motif exhibited significantly reduced expression in *PCIF1* knockout cells. In contrast, transcripts that use the VA_+1_RR TSS motif were largely unaffected. Notably, transcripts normally have small differences in methylation (i.e., methylation level at 0.9 vs 0.98). Thus, m^6^Am is unlikely to be a general stabilization mark in mRNA since it only affects transcripts based on promoter sequences. Instead, these different stoichiometries of m^6^Am are likely to be the consequence of different transcription mechanisms. Thus, it will be important to assess how these different transcription mechanisms use m^6^Am for gene expression.

PCIF1 is known to be associated with RNA polymerase II and is recruited to promoter regions during transcription^28^. Thus, PCIF1 is ideally positioned to regulate transcription processes. It is interesting to speculate that m^6^Am may provide a mark that enhances subsequent elongation and thus maintains high overall expression for transcription initiation complexes that assemble on the SSCA_+1_GC TSS motif. Other transcription initiation complexes, such as those using the VA_+1_RR TSS motif, may not need this mechanism. However, our data cannot provide further details on whether the loss of m^6^Am is related to exact mechanisms such as transcription initiation selection, elongation, or premature termination. Notably, the recent study by An et al.^39^ suggested that the loss of m^6^Am is related to enhanced premature termination and therefore leads to reduced RNA 5’ end expression. An et al. proposed that m^6^Am can sequester PCF11, a m^6^Am reader, and thereby promote transcription by reducing premature transcription termination^39^. However, it is still unclear whether the transcripts from the SSCA_+1_GC TSS motif are indeed more preferentially bound by PCF11. To better understand how PCIF1 regulates transcription, assays with transcription-start nucleotide resolution will be required.

Although m^6^Am and m^6^A are chemically similar, these two modifications appear to have very different biological functions. It is well known that m^6^A is a mark for RNA instability through the recruitment of YTHDF proteins^40^. However, we find no correlation between m^6^Am and RNA instability. Additionally, our previous YTHDF1, YTHDF2, and YTHDF3 iCLIP studies did not show binding at mRNAs 5’ ends^41^, which suggests that YTHDF proteins do not bind m^6^Am. Thus, specific m^6^Am-binding proteins might enable its unique functions in transcription.

Although most studies of m^6^Am and PCIF1 focus on mRNAs, we find that m^6^Am in snRNAs exhibit substantially higher variability and regulation than that in mRNA. Early biochemical studies of snRNA composition demonstrated that the first nucleotide was generally Am in all Pol II-derived snRNAs^10^. CROWN-seq generally supports this finding since most snRNAs have low m^6^Am stoichiometry. However, the previous study mainly focused on HEK293T cells^10^, which have very low m^6^Am in snRNA. In this study, nine different cell types were sequenced. These cell lines showed highly variable m^6^Am in snRNA. In some cases, several snRNAs can reach m^6^Am stoichiometry up to 0.70-0.89. These data raise the possibility that m^6^Am may affect snRNA functions, such as splicing and gene transcription^12,42^, and *PCIF1* knockout phenotypes may be due to altered snRNA.

Notably, m^6^Am in snRNA is highly regulated by FTO, which is consistent with our earlier findings^10^. However, the previous study did not have transcript isoform level resolution in analyzing the effect of FTO demethylation. With CROWN-seq, we find that FTO has markedly different effects on different snRNAs, where some snRNAs appear highly demethylated by FTO while others are insensitive to FTO. Some snoRNA, and snRNA/snoRNA pseudogenes are also demethylated by FTO. Notably, FTO depletion affects numerous aspects of cell function^43^. Our results thus raise the possibility that FTO-depletion phenotypes may result from increased m^6^Am levels in snRNAs, snoRNAs, or their pseudogenes.

### Limitations of the study

One limitation of CROWN-seq is that it can be difficult to align sequencing reads to the genome. Unlike normal reads, which contain A, G, C, and U, most reads in CROWN-seq comprise only G, C, and U due to the conversion of A’s. This makes it difficult to align reads to highly similar genes, such as snRNA isoforms and pseudogenes which have very similar 5’ ends. For this reason, only a small fraction of reads from snRNA and pseudogenes were uniquely mapped to one genomic location and were used in this analysis. To better understand m^6^Am in these 5’ ends with similar sequences, future optimization is desired to increase the read lengths, which can help distinguish these sequences from each other. This requires technical innovations in reducing RNA fragmentation during sodium nitrite conversion, which comes from acid-catalyzed depurination and backbone cleavage^21^.

In this study, we quantified m^6^Am in nine different cell lines, which cover a wide range of PCIF1 expression levels. Although we found high m^6^Am stoichiometries in all cell lines, it is possible that some cells or tissues have more variable m^6^Am levels. In our previous study, mass spectrometry showed that the CCRF-SB cell line has relatively low m^6^Am stoichiometry at ∼67.6%^1^. However, these cells exhibit very slow growth as reported previously^1^. As a result, we were unable to obtain enough amounts of mRNA needed for CROWN-seq. Future CROWN-seq studies may lead to the identification of cell types or contexts with dynamic m^6^Am landscapes.

The last limitation of this study is that the focus of this study was to quantify m^6^Am and to predict potential functions using *PCIF1* knockout cells. However, it is possible that PCIF1 has non-catalytic functions that may contribute to the *PCIF1* knockout phenotype. Future experiments using catalytic-dead PCIF1 can be useful to distinguish between the catalytic and non-catalytic functions of PCIF1.

## ACKNOWLEDGEMENTS

We thank members of the Jaffrey lab for their comments and suggestions throughout the duration of this project. We thank members of the Genomics core facility at Weill Cornell Medicine. This work is supported by NIH grants R35 NS111631, S10 OD030335, RM1 HG011563, and MH121072 (S.R.J.).

## AUTHOR CONTRIBUTIONS

J.F.L., B.R.H., and S.R.J. designed the experiments. J.F.L., B.R.H., and L.N. performed experiments. J.F.L. and B.R.H. analyzed experimental data. J.F.L and S.R.J. wrote the manuscript.

## DECLARATION OF INTERESTS

S.R.J is a scientific founder of, advisor to, and/or owns equity in Chimerna Therapeutics, Lucerna Technologies, and 858 Therapeutics.

## SUPPLEMENTAL TABLES

**Table S1.** Comparison of m^6^Am mapping methods.

**Table S2.** The design of m^6^Am standards.

**Table S3.** mRNA m^6^Am stoichiometry in different cell lines.

**Table S4.** Gene Ontology enrichment of genes with relatively low m^6^Am sites.

**Table S5.** Comparing A-transcription-start nucleotide expression between wild-type and *PCIF1* knockout cells.

**Table S6.** m^6^Am stoichiometry of A-TSN in snRNA, snoRNA, and pseudogenes.

**Table S7.** Oligos used in this study.

## METHODS

### EXPERIMENTAL MODEL AND SUBJECT DETAILS

#### Cell lines

HEK293T (wild-type, *PCIF1* knockout, and FTO knockout cells), A549 (wild-type and *PCIF1* knockout), HCT-116, Huh-7, and HT-29 cells were maintained in DMEM (Gibco #11995065). HepG2 and CCD841 CoN cells were maintained in EMEM (ATCC #30-2003). K562 and Jurkat E6.1 cells were maintained in RPMI1640 (Gibco #11875093). All media was supplemented with 10% FBS and 1X penicillin-streptomycin (Gibco #15140148). Cells were grown at 37 °C with 5% CO_2_. We followed the instructions from the manufacturer to maintain the cells.

### METHODS DETAILS

#### RNA extraction and mRNA purification

Cellular total RNA in TRIzol LS (ThermoFisher #10296028) was extracted by Direct-zol RNA Miniprep kit (Zymo #R2070) or by Phenol Chloroform extraction. mRNA was purified by NEBNext Oligo d(T)25 Beads (NEB #E7499) or Dynabeads Oligo (dT)25 (Ambion #61002) based on mRNA purification from total RNA protocol of Dynabeads Oligo (dT)25 (Ambion #61002).

#### m^6^Am standard preparation

We used *in vitro* transcription to prepare m^7^G capped m^6^Am- or Am-initiated transcripts, which are based on HiScribe® T7 mRNA Kit with CleanCap® Reagent AG (NEB #E2080S). We first obtained DNA templates from IDT gBlock. In total, five DNA templates which are identical expect for the 6-nt long barcode 42-nt downstream to the TSS were used (**Table S2**). The DNA templates contain 5’-TAATACGACTCACTATAAG-3’ T7 promoter for *in vitro* transcription. We used CleanCap® Reagent AG (3’ OMe) (TriLink #N-7413), which is included in NEB #E2080S, to generate m^7^G-ppp-Am modified transcripts. We used CleanCap® Reagent M6 (TriLink #N-7453) to generate the m^7^G-ppp-m^6^Am modified transcripts. The RNAs made by *in vitro* transcription were DNase I treated, purified, and then quantified by both Agilent TapeStation (RNA high sensitivity assay). We then mixed the Am and m^6^Am modified oligos to generate m^6^Am standards with expected m^6^Am stoichiometry at 0%, 25%, 50%, 75%, and 100% m^6^Am stoichiometry. Notably, the guaranteed purity of the CleanCap® Reagent M6 is >95%. The CleanCap® Reagent M6 can contain m^7^G-ppp-AmG analog, which results in the reduced non-conversion rate in CROWN-seq.

#### Genomic assembly and annotations

The genomic sequence and annotations of Gencode v45 primary assembly were used in this study.

#### GLORI experiment

To validate whether sodium nitrite conversion can convert Am into Im, we spiked ∼0.01 ng Am transcripts (ERCC-00057-1-TCGTCG) into ∼250 ng poly(A) selected mRNA for GLORI assay. Ligation-based GLORI protocol was used in this study. Notably, the Am transcripts were decapped by mRNA Decapping Enzyme (NEB #M0608S) in advance. We first fragmentized the input RNA into ∼200 nt long fragments (NEBNext Magnesium RNA Fragmentation Module (NEB #E6150S), 94°C, 2 minutes). The fragmentized RNAs were then A-to-I converted based on the GLORI protocol^20^: we converted the glyoxal-protected RNA by 750 mM NaNO_2_ at 16°C for 8 hours and 4°C overnight. The RNA was then deprecated in a deprotection buffer at 95°C for 10 minutes. The deprotected RNA was then T4 PNK (NEB #M0210S) treated and processed to ligation-based small RNA-seq library preparation^46^. Notably, the 5’ adapter in library preparation contains an 11 nt UMI sequence (see **Table S7**).

#### GLORI data processing

GLORI libraries were analyzed based on a modified mRNA bisulfite sequencing pipeline^47^. The first 10 bases in GLORI libraries made with eCLIP protocol were first extracted by a customized script. GLORI reads were first quality trimmed by Cutadapt^48^. For the GLORI library generated by eCLIP protocol, the parameters are --max-n 0 --trimmed-only -a AGATCGGAAGAGCGTCGTG -e 0.1 -q 30 -m 40 --trim-n; for GLORI library prepared by ligation based protocol generated in this study, the parameters are -m 32 -j 4 -q 20 -e 0.25 -a AGATCGGAAGAGCACACGTC -A ATATNNNNNNNNNNNAGATCGGAAGAGCGTCGTG. After pre-processing, the reads were firstly A-to-G converted and aligned to A-to-G (positive strand) and T-to-C (negative strand) converted reference genome and transcriptome by Hisat2-2.1.0^49^. Parameters in alignment: -k 5 –fr –rna-strandness FR –no-temp-splicesite –no-mixed. Only unique alignments were used. After alignment, the base information in sequences was restored so that m^6^Am signals can be reflected by the A-to-G mismatches. No further transcriptome alignment was performed on the unmapped reads. After alignment, a customized script based on Pysam^50^ was used to pileup every single base to obtain the A, C, G, and U counts. Every single base was assigned to a transcript isoform if possible (order: mRNA > lncRNA > functional RNAs > pseudogenes). Non-conversion rate is defined as the number of A counts against the sum of A count and G count. Filters were applied to obtain high-quality non-converted A (m^6^A/m^6^Am) signals in a gene-specific manner: (1) only genes with at least 1000 counts were analyzed; (2) gene-specific non-conversion rates were computed for Binomial test on the frequency of non-conversion and sites with Binomial test P-value < 0.05 were used; (3) reads with more than 3 non-converted As were considered as noise and discarded; (4) sites with more than 5% signals were discarded due to the site may fall in a conversion-resistant region; (5) Only sites with no less than 20 reads covered and non-conversion rates over 0.1 were considered as m^6^A/m^6^Am sites. (6) Non-conversion rates of the same site from different replicates were averaged. Details of this pipeline can be found at https://github.com/jhfoxliu/GLORI_pipeline.

#### ReCappable-seq library preparation

A modified ReCappable-seq protocol^18^ was developed to reduce background, reduce material loss, and increase the utility of mapped reads. Several steps of library construction are now performed while the 5’ desthiobiotinylated cap is bound to streptavidin beads. This reduces the opportunity for carry-through of random fragmentation products to occur that would previously result in non-cap-derived 5’ ends to be ligated. Next, 5’ adapters with unique molecular indexes (UMIs) are used to permit robust PCR duplicate removal. Finally, ∼160 spike-in mRNAs from SIRV-ERCC Spike-in mixture (Lexogen #051.03) with single defined 5’ termini are used, which are used during analysis to build a dynamic thresholding pipeline that exclude false positive start sites. A complete step-by-step protocol as performed here will be available on the accompanying GitHub page (see **Data and Code Availability**).

5 μg total RNA was used as input for all experiments. RNA was denatured at 65 °C for 2 minutes before reaction mixes were added. First, 5’-phosphorylated RNAs were dephosphorylated using 25 U Quick CIP (NEB #M0525L) in a 50 μL reaction for 30 minutes at 37 °C. The reaction was cleaned using a Zymo RCC-5 column following the manufacturer’s >200 nt protocol and eluted with 20 μL water. m^7^G capped RNAs were then decapped using 200 U yDcpS (NEB #M0463S) for 1 hour at 37 °C. This unique decapping enzyme liberates m^7^GMP, resulting in mRNAs with a 5’-diphosphate. The reaction was cleaned and eluted as before. Next, the 5’-diphosphorylated mRNAs were recapped with desthiobiotin-GTP (DTB-GTP) using vaccinia capping enzyme (5 μL VCE buffer, 0.5 μL inorganic pyrophosphatase (NEB #M0361S), 5 μL DTB-GTP (5 mM; NEB #N0761S), 50 U VCE (#M2080S)) for 45 minutes at 37 °C. The reaction was clean as before, however, a total of 4 washes were performed to ensure the complete removal of excess DTB-GTP. RNA was then fragmented by incubating at 95 °C for 2.5 minutes in a 25 μL reaction containing 100 mM Tris-HCl pH 8.0 and 2 mM MgCl_2_. Fragmented RNA was placed on ice and brought to 30 μL with water. Streptavidin beads (NEB #S1421S) were washed in a high salt wash buffer (10 mM Tris-HCl pH 7.5, 2 M NaCl, 1 mM EDTA) and resuspended in the high salt buffer at 4 mg/mL. 30 μL beads were added to 30 μL fragmented RNA and incubated for 45 minutes at room temperature with agitation. Beads were washed twice in a high salt buffer, twice in a lower salt buffer (10 mM Tris-HCl pH 7.5, 250 mM NaCl, 1 mM EDTA), and twice in PNK wash buffer (20 mM Tris-HCl pH 7.5, 10 mM MgCl_2_, 0.2% Tween). Beads were next resuspended in 40 μL PNK reaction mix (8 μL 5X pH 6.5 PNK buffer (350 mM Tris-HCl pH 6.5, 50 mM MgCl_2_, 5 mM DTT), 1 μL T4 PNK (NEB #M0201S), 1 μL

RNaseOUT (ThermoFisher #10777019) and incubated at 37 °C for 30 minutes with agitation to remove 3’ phosphates resulting from the fragmentation. Beads were washed once in PNK wash, once in the high salt wash, then twice again in PNK wash. Next, a 3’ adapter (see **Table S7**) was added to RNA by resuspending beads in 40 μL 3’ ligation reaction mix (4 μL T4 RNA ligase buffer, 2 μL T4 RNA ligase 2 truncated KQ (NEB #M0373L), 1 μL RNaseOUT, 2 μL L7 adapter (20 μM stock, see **Table S7**), 16 μL of 50% PEG-8000) and incubated at 25 °C for 2 hours. The beads were washed once in PNK wash, once in high salt wash, twice in lower salt wash, then resuspended in 30 μL lower salt wash containing 1 mM biotin (ThermoFisher #B20656) to elute DTB-capped RNA fragments. The eluted RNA was cleaned by ethanol-AMPure XP (1.8 volumes AMPure XP, then 1.5 volumes 100% ethanol). To increase stringency, the streptavidin bead enrichment was repeated omitting enzymatic steps and instead washing three times with high salt and then three times with lower salt wash, and the eluate was cleaned again by ethanol-AMPure XP. The DTB-GTP cap was removed using 0.5 U/µl RppH (NEB #M0356S) in 1X ThermoPol buffer (NEB #M0356S) and incubating at 37 °C for 1 hour. The resulting 5’-monophosphate RNA fragments were purified by ethanol-AMPure XP. 30 pmol of a 5’ adapter was ligated for 3 hours at 25 °C with 2 U/μL T4 RNA ligase 1 (NEB #M0437M). This RNA adapter contains an 11 nt UMI followed by a fixed sequence (AUAU) at its 3’ end. The UMI allows robust duplicate removal, and the fixed sequence provides an anchor point to correctly identify the first nucleotide of the mRNA. The ligation reaction was inactivated by heating at 65°C for 10 minutes and then immediately used in a reverse transcription reaction. 3 pmol of ReCappable-seq RT primer was annealed to the 3’ adapter of RNA fragments by heating to 65°C for 5 minutes and cooling to 25 °C at a rate of 0.1 °C/sec. Reverse transcription was carried out at 55 °C for 45 minutes in a reaction containing 0.5 mM dNTPs, 5 mM DTT, 20 U RNaseOUT, 50 mM Tris-HCl pH 8.3, 75 mM KCl, and 300 U SuperScript III (ThermoFisher #18080044). Following heat inactivation, the reaction was cleaned using ethanol-AMPure XP and cDNA was resuspended in 21 μL. The final PCR was performed using 8 μL cDNA in a 40 μL reaction containing 1X Phusion HF master mix (NEB #M0531L) and 4 μL each of a unique i5 and i7 barcoded primer combination for each sample (NEB #E7600S). Cycling conditions were typically 98 °C 2 minutes, then 11-13 cycles of 98 °C 15 seconds, 65 °C 30 seconds, 72 °C 30 seconds, with a final 5 minute 72 °C extension. The optimal number of cycles for each library was determined by performing a set of test cycles using 1 μL cDNA in a 20 μL reaction. PCR libraries were purified with 2 rounds of bead clean-up using 0.9X volume SPRIselect beads. Libraries were pooled at equimolar concentrations and sequenced in paired-end mode with 50-150 bp reads depending on the library on either an Illumina NovaSeq, NextSeq, or HiSeq (please refer to GEO accession for specific details for each library).

#### ReCappable-seq analysis

The beginning of read 1 is the UMI plus an ATAT spacer sequence, and the nucleotide directly following this is the TSS. Reads were first filtered to identify pairs with the correct UMI+ATAT sequence, then the UMI was added to FASTQ headers using UMI-tools v1.1.1^51^. ATAT sequence discarded. Adapters were trimmed using Cutadapt v3.4^48^. Next, reads mapping to ribosomal RNA and small non-coding RNAs were filtered away by aligning to these sequences using bowtie2 v2.4.2^52^. Reads were then aligned to GRCh38 and m^6^Am standard sequences using HISAT2^49^. The alignment results were deduplicated by UMI-tools (--paired --chimeric-pairs=discard --unpaired-reads=discard --method=unique). Only reads without 5’ softclipping were used. A customized script based on Pysam^50^ was used to extract the 5’ ends from the BAM file. To annotate the sites by a gene, the 5’ ends were firstly annotated by the nearest TSS within the 100 bp region. If multiple annotations were found, the annotation was selected by the priority of snRNA > snoRNA > mRNA > lncRNA > others. BEDtools v2.27.1^53^ was used to find the nearest annotation. To more accurately estimate the expression levels of each TSN, we normalized the read counts using the “RUVg” function in RUVSeq pacakge^54^. To calculate the expression levels of TSNs in wide-type cells, we calculated the TPM values based on the normalized read counts. To calculate the differential expression between wild-type and *PCIF1* knockout cells, the normalized read counts were proceeded by DESeq2^45^.

#### CROWN-seq library preparation

CROWN-seq uses the glyoxal-based guanosine protection protocol from GLORI^20^ and a TSN enrichment protocol that is modified from ReCappable-seq^18^. In CROWN-seq, glyoxal protection is very important to prevent both internal G’s from being converted into xanthosine, which can interrupt base pairing and cause mutations during reverse transcription^55^. Because N7-methyl does not interrupt the interaction between glyoxal and N1 and N2 positions of guanosines, glyoxal protection is also very useful to prevent m^7^G from being converted, which can help 5’ end enrichment. After glyoxal protection, A bases are deaminated into inosines by sodium nitrite.

After deamination, the 5’ end RNA fragments with a m^7^G cap were enriched by ReCappable-seq workflow, where the m^7^G caps were replaced by a 5’ desthio-biotinylated cap for enrichment by streptavidin beads. 3’ adapter and 5’ adapter (with unique molecular indexes (UMIs)) were ligated to the enriched 5’ RNA fragments, so that the library can be made by reverse transcription followed by indexing PCR. Detailed workflow is described below.

Conversion. 0.8-2.5 μg oligo(dT) selected RNA was used as input. RNA was first diluted in 14 μl water. To perform glyoxal protection, 6 µl 8.8 M glyoxal and 20 μl DMSO were then added to the diluted RNA and well mixed. The 40 µl mix was first incubated at 50 °C for 30 minutes, then 10 μl boric acid was added to the mix. The 50 µl mix was then incubated for an additional 30 minutes at 50 °C. After protection, the 50 μl protected RNA was mixed with 50 μl deamination buffer (25 μl 1500 mM NaNO_2_, 4 μl 500 mM MES, pH 6.0, 10 μl 8.8 M glyoxal, and 11 μl water). The deamination reaction was performed at 16 °C for 8 hours. After deamination, the RNA was recovered by ethanol precipitation. To remove the glyoxal adduct from the RNA, the RNA pallet was dissolved in 50 μl deprotection buffer (500 mM TEAA pH=8.6, 47.5% deionized formamide) and was incubated at 95 °C for 10 minutes. After incubation, the reaction was brought to 250 μl with water. Converted RNA was purified by ethanol precipitation and eluted in 39 μl water for 5’ end enrichment. The converted RNA was stored at -80 °C before 5’ end enrichment.

Recapping. To eliminate the contamination of RNA with 5’-triphosphate and 5’-monophosphate, 5 µl 10X CutSmart buffer, 5 µl Quick CIP (5 U/µl) (NEB #M0525L), and 1 µl SUPERase·In RNase inhibitor (ThermoFisher #AM2696) were added to the 39 μl converted RNA to set up a dephosphorylation reaction. The dephosphorylation reaction was performed at 37 °C for 30 minutes. The reaction was cleaned up using Zymo RCC-5 column and the RNA was eluted in 42 µl water. To decap the m^7^G capped RNA, a 50 µl decapping reaction was set up by adding 5 µl 10X yDcpS buffer, 2 µl (200 U) yDcpS (NEB #M0463S), and 1 µl SUPERase·In to the 42 µl dephosphorylated RNA. The decapping reaction was performed at 37 °C for 1 hour. This unique decapping enzyme liberates m^7^GMP, resulting in mRNAs with a 5’-diphosphate. The reaction was cleaned and eluted as before. The reaction was cleaned up using Zymo RCC-5 column and the RNA was eluted in 33.5 µl water. The 5’-diphosphorylated mRNAs were recapped with desthiobiotin-GTP (DTB-GTP, NEB #N0761S) using vaccinia capping enzyme (VCE, NEB #M2080S) (5 μL VCE buffer, 0.5 μL inorganic pyrophosphatase (NEB #M0361S), 5 μL DTB-GTP (5 mM), 50 U VCE, 1 µl SUPERase·In) at 37 °C for 1 hour. The reaction was cleaned up using Zymo RCC-5 column and the RNA was eluted in 30 µl water. Now the RNA is ready for streptavidin enrichment.

5’ enrichment. Streptavidin beads (NEB #S1421S) were washed in a high salt wash buffer (10 mM Tris-HCl pH 7.5, 2 M NaCl, 1 mM EDTA) and resuspended in the high salt buffer at 4 mg/mL. To enrich the RNA and tag the 5’ and 3’ end by the specific adapter, the 30 µl recapped RNA was first mixed with 30 µl streptavidin beads and incubated at room temperature for 45 minutes with agitation. Beads were washed twice in high salt buffer, twice in a lower salt buffer (10 mM Tris-HCl pH 7.5, 250 mM NaCl, 1 mM EDTA), and twice in PNK wash buffer (20 mM Tris-HCl pH 7.5, 10 mM MgCl_2_, 0.2% Tween). To remove 3’ phosphates resulting from fragmentation during conversion, beads were resuspended in 50 µl PNK reaction without ATP (5 µl 10X PNK buffer, 1 µl T4 PNK (#M0201S), 1 µl SUPERase·In, 43 µl water), and incubate at 37 °C for 30 minutes with agitation. The beads were then washed once in PNK wash buffer, once in 2 M NaCl wash, and twice in PNK wash. Next, RNA was ligated to a 74 nt-long 3’ adapter (**see Table S7**) in the following 40 μl 3’ ligation mix: 4 μL T4 RNA ligase buffer, 2 μl T4 RNA ligase 2 truncated KQ (NEB #M0373L), 1 µl SUPERase·In, 2 μl extended-L7 adapter (20 μM stock), 16 μl of 50% PEG-8000) and incubated at 25 °C for 2 hours. After incubation, the reaction buffer was removed by washing once with high salt buffer and twice with PNK wash buffer. To remove the exceeded adapter, the beads were incubated in 50 μl adapter digestion reaction (40 μl water, 5 µl 10X RNA ligase buffer, 1 µl RecJf (NEB #M0264S), 1 5’ Deadenylase (NEB #M0331S), 1 µl SUPERase·In) at 30 °C for 15 minutes then at 37 °C for 15 minutes. The beads were washed once with PNK wash, once with high salt buffer, and twice with low salt buffer. To elute the DTB labeled RNA, beads were then suspended with 30 µl low salt wash buffer containing 1 mM free D-biotin (ThermoFisher #B20656) and incubate at room temperature for 1 hour. The DTB-labeled RNA was purified with ethanol-AMPure XP (RNA:beads:ethanol=1:2:3) and eluted in 30 µl water. To increase stringency, the streptavidin bead enrichment was repeated omitting enzymatic steps and instead washing three times with high salt and then three times with lower salt wash, and the eluate was cleaned again by ethanol-AMPure XP.

5’ adapter addition. The DTB-GTP cap was removed using 0.5 U/µl RppH (NEB #M0356S) in 1X ThermoPol buffer (NEB #M0356S) and incubating at 37 °C for 1 hour. The resulting 5’-monophosphate RNA fragments were purified by ethanol-AMPure XP and eluted in 10 μl water. 1 μl (10 pmol) reverse transcription primer (see **Table S7**) was pre-annealed to the templates by heating up to 75 °C for 5 minutes, then 37 °C for 15 minutes, 25 °C for 15 minutes, and chilled at 4 °C. 10 pmol of a 5’ adapter (see **Table S7**) was ligated for 3 hours at 25 °C with 2 U/μl T4 RNA ligase 1 (NEB #M0437M). This RNA adapter contains an 8 nt- or 11 nt-long UMI followed by a fixed sequence (AUAU) at its 3’ end. The UMI allows robust duplicate removal, and the fixed sequence provides an anchor point to correctly identify the first nucleotide of the mRNA. 40 μl ligation product was used.

cDNA synthesis and PCR. Reverse transcription was carried out at 50 °C for 45 minutes in a 50 μl reaction containing 0.5 mM dNTPs, 5 mM DTT, 20 U RNaseOUT, 50 mM Tris-HCl pH 8.3, 75 mM KCl, and 300 U SuperScript III. To perform indexing PCR, 40 μl Phusion master mix (NEB # M0532L) was added to the reverse transcription product, along with 5 μl i5 indexing primer and 5 μl i7 indexing primer (NEB #E7600S). Cycling conditions were typically 98 °C 2 minutes, then 16 cycles of 98 °C 15 seconds, 65 °C 30 seconds, 72 °C 30 seconds, with a final 5 minute 72°C extension. Two rounds of 0.9X AMPureXP bead purifications were performed to remove primers. Normally ∼10 ng indexed library was obtained for each library. The libraries were mixed and sequenced by NovaSeq 6000 or NovaSeqX.

#### CROWN-seq data processing

The read pairs were firstly quality trimmed by Cutadapt^48^: -m 32 -q 20 -e 0.25 -a AGATCGGAAGAGCACACGTC. For the 8 nt-long 5’ adapter, -A ATATNNNNNNNNAGATCGGAAGAGCGTCGTG was used; for the 11 nt-long adapter, -A ATATNNNNNNNNNNNAGATCGGAAGAGCGTCGTG was used. Then the UMI along with the fixed ATAT spacer sequences were extracted by UMI-tools^51^. The alignment process was modified from the previous RNA bisulfite alignment strategy^47^. In brief, *in silico* converted read pairs (read1 A-to-G, read2 T-to-C) were aligned by HISAT2^49^ against A-to-G converted (for positive strand) and T-to-C converted (for negative strand) reference genome and transcriptome first (key options: -k 5 –fr –rna-strandness FR –no-temp-splicesite –no-mixed). Then the unique alignments were extracted and the *in silico* converted reads were inverse-transformed to the original format. Since two sequences after conversion can be easily confused, we require the best alignment results can be well distinguished from the secondary alignments. Here, the alignment scores (AS tag in Hisat2 alignments, higher is better) of the best alignments should be higher than -10. Meanwhile, the difference between the best alignments and secondary alignments should be larger than 9. For paired-end alignments, the alignment scores of read1 and read2 were summed. Only read1 was used in the 5’ end analysis. Only read1 reads without 5’ end softclips were used. Pileup was performed to obtain the read coverages of every 5’ end in the transcriptome. Non-conversion rates of the transcription start nucleotides were calculated by A counts over A and G counts.

To annotate the TSNs mapped in CROWN-seq, we used the TSSs in Gencode v45 as the reference TSS positions. We first calculated the distance between the mapped TSNs and the annotated TSSs by BEDtools^53^. We then tried to assign a TSN to a gene if there was an annotated TSS <100 nt away. Because there can be multiple annotations available, we used the following priority in selecting gene annotations: snRNA > snoRNA > protein-coding > lncRNA > others. We also annotated TSNs which come from RNA highly similar to snRNA, snoRNA, or their pseudogenes. To do so, we first built a BLASTn database containing all snRNA, snoRNA, and their pseudogene sequences from Gencode v45. We then performed BLASTn (BLAST 2.9.0+^56^) on the A-TSNs along with the first 50 nt downstream sequences to examine the similarity to the known snRNA, snoRNA, and pseudogenes. The following parameter was used: -qcov_hsp_perc 50 -perc_identity 50 -word_size 10. Sequences with bitscore ≥50 were considered as snRNA/snoRNA-like. We also annotated uORF and IRES elements based on ORFdb^57^ and IRES atlas^58^, respectively. The related pipeline and scripts are available at https://github.com/jhfoxliu/CROWN-seq.

#### RT-qPCR

1 μg total RNA was used as input. The RNA was then mixed with 1 μl Oligo dT(18) (100 pmoles) (ThermoFisher #SO131), and 1 μl dNTP in 14.5 μl total volume. The mix was incubated at 65 °C for 5 minutes, then on ice for >30 seconds. After the incubation, 4 μl 5X RT mix (Maxima H-buffer, ThermoFisher #EP0751), 0.5 μl RNaseOUT (ThermoFisher #10777019), and 1 μl Maxima H-RTase were added to the mix. Reverse transcription was performed at 25 °C for 10 minutes, then 50 °C for 30 minutes. After reverse transcription, 1 μl cDNA was used for qPCR. In addition to the cDNA input, the qPCR buffer contains 10 μl Power SYBR Green PCR Master Mix (ThermoFisher #368577), 0.5 μl forward primer, 0.5 μl reverse primer, and 8 μl water. qPCR was performed based on the standard quantification program in QuantStudio™ 5 System.

#### Gene ontology analysis

Gene ontology analyses were performed with R package ClusterProfiler^59^. P-value cutoffs were set to 0.05 and q-value cutoffs were set to 0.1. “Cellular Components” and “Biological Process” terms were analyzed. Importantly, corresponding gene sets, rather than all genes, were used as the backgrounds in term enrichment computation. Since the output terms were normally redundant, terms were de-redundancy by the “simplify” function in R package GOSemSim^60^ (cutoff=0.7, by=“p.adjust”, select_fun=min).

#### Motif analysis

To search for the potential motifs for elements related to transcription initiation, we used “re” package in Python to match specific motifs, which are indicated in the figure legends.

To search for the transcription factor binding sites, we used FIMO^61^ to scan for motifs in HOCOMOCO v11 core motifs database^62^.

#### RNA secondary prediction and minimal free energy calculation

ViennaRNA package (version 2.5.1) was used to perform RNA secondary structure prediction^63^. The RNAfold Python API “RNA” was used in the analysis. The folding temperature was set to 37°C. The minimum free energy of the predicted structure was used.

### QUANTIFICATION AND STATISTICAL ANALYSIS

Quantitative and statistical methods are described above and in figure legends according to their respective technologies and analytic approaches. Statistical analysis and visualization were mainly performed with Python (version 3.8.7). R (version 4.2.2) was used in differential gene expression analysis and Gene Ontology analysis.

Versions of key Python packages: numpy (1.23.5); pandas (1.5.2); scipy (1.9.3); matplotlib (3.6.2); seaborn (0.12.1); matplotlib-venn (0.11.9).

Versions of key R packages: DESeq2 (1.38.1); clusterProfiler (4.6.0); enrichplot (1.18.3); GOSemSim (2.24.0); org.Hs.eg.db (3.16.0).

All boxplots and violin plot summary statistics show the median and IQR of the underlying data. Statistical tests are described in the appropriate figure legends. Student’s t-test was applied for two sample non-paired comparisons. One-sided or two-sided testing was performed according to figure legends. If possible, we omitted significance “stars” from figures; p-values (or equivalent) are instead reported.

### DATA AND CODE AVAILABILITY

All sequencing data can be accessed from NCBI Gene Expression Omnibus: GSE188510 for ReCappable-seq; GSE233655 for CROWN-seq data.

Because of the size limit of supplemental tables, several large tables (e.g., the table for all mRNA m^6^Am sites quantified in this study) were not directly included in the **Supplemental**

**Tables**. These large supplemental tables have been uploaded to Zenodo: https://zenodo.org/records/12760731 (DOI: 10.5281/zenodo.12760731).

All original code has been deposited on GitHub:

- ReCappable-seq analysis: https://github.com/jhfoxliu/ReCappable-seq
- GLORI analysis: https://github.com/jhfoxliu/GLORI_pipeline
- CROWN-seq analysis: https://github.com/jhfoxliu/CROWN-seq

Any additional information required to reanalyze the data reported in this paper is available from the lead contact upon request.

## MATERIALS AVAILABILITY

This study did not generate new unique reagents.

## Notes

https://www.ncbi.nlm.nih.gov/geo/query/acc.cgi?acc=GSE233655

https://www.ncbi.nlm.nih.gov/geo/query/acc.cgi?acc=GSE188510

https://zenodo.org/records/12760731

### Competing Interest Statement

S.R.J. is the co-founder and/or has equity in Chimerna Therapeutics, 858 Therapeutics, and Lucerna Technologies.

### Summary of Updates

Manuscript file not changed. Update supplmentary file.

