## Supplementary material for "Decoding m^6^Am by simultaneous transcription-start mapping and methylation quantification": Supplmentary figures and notes

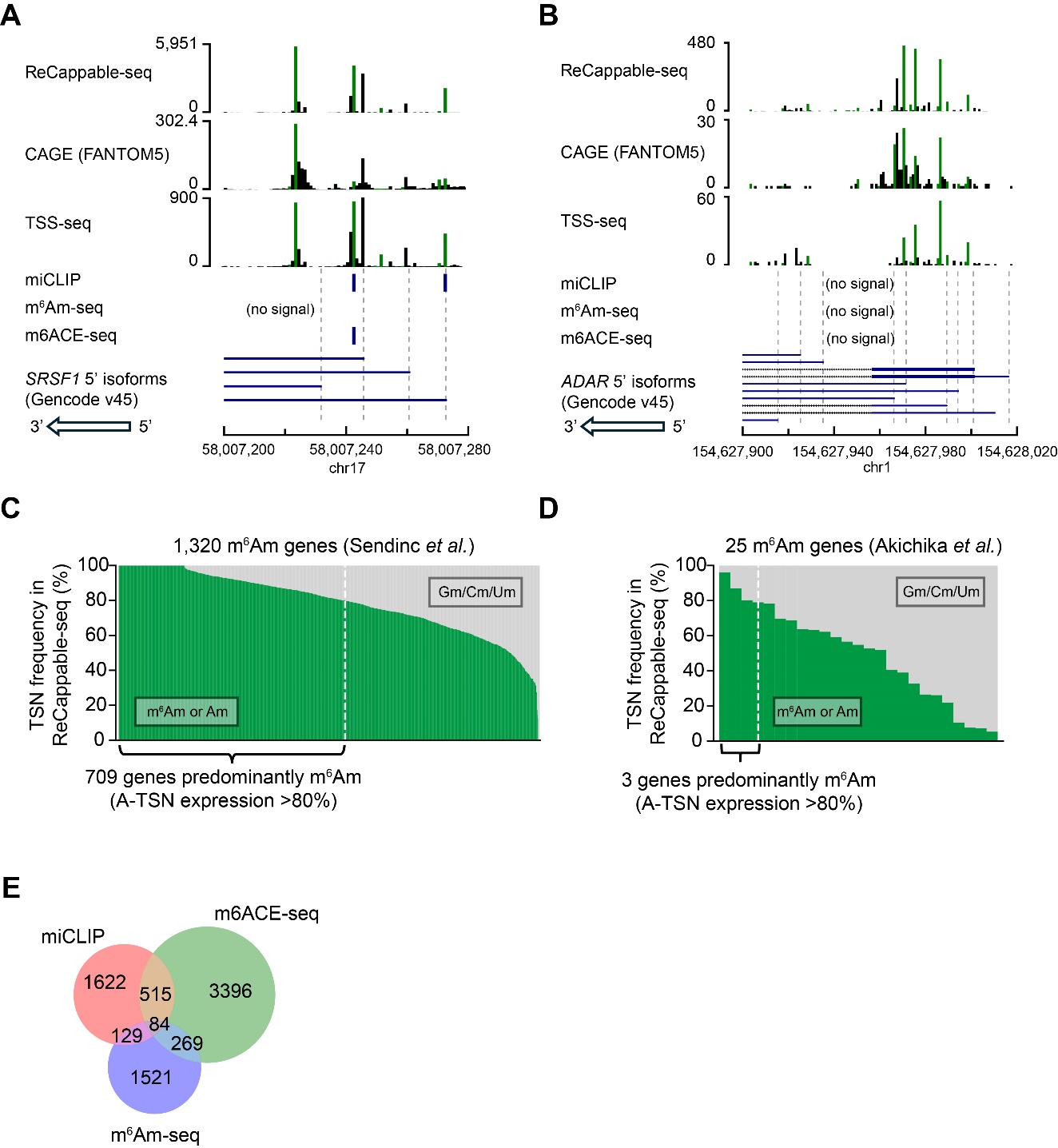

**Figure S1. Corresponding to Figure 1.**

**(A)** and **(B)**. Corresponding to **Figure 1B** and **1C**, the heterogeneity of TSS usage in *SRSF1* and *ADAR* can also be observed in TSS mapping data generated by CAGE and TSS-seq. In these plots, HEK293T CAGE data were downloaded from FANTOM5^1^; HEK293T TSS-seq data were generated in our previous study^2^.

**(C)** and **(D).** Similar to Figure 1B, shown are the m^6^Am genes classified by **(C)** Sendinc et al. ^3^ and **(D)** Akichika et al. ^4^. For each gene, the percentage of transcripts starting with m^6^Am/Am (in green) or Gm/Cm/Um (in gray) are shown. The transcription-start nucleotide frequencies are obtained by ReCappable-seq data of HEK293T cells. For **(D)**, the 25 m^6^Am genes are collected from the supplemental table “List of genes whose TEs are up- or down-regulated upon CAPAM KO” in the original paper.

**(E)** Venn diagram showing the inconsistency between m^6^Am maps generated by the existing m^6^Am mapping methods. In this analysis, only m^6^Am sites mapped to the primary chromosome are used. Source of data: miCLIP, Boulias et al. ^5^; m^6^Am-seq, Sun et al. ^6^; m6ACE-seq, Koh et al. ^7^.

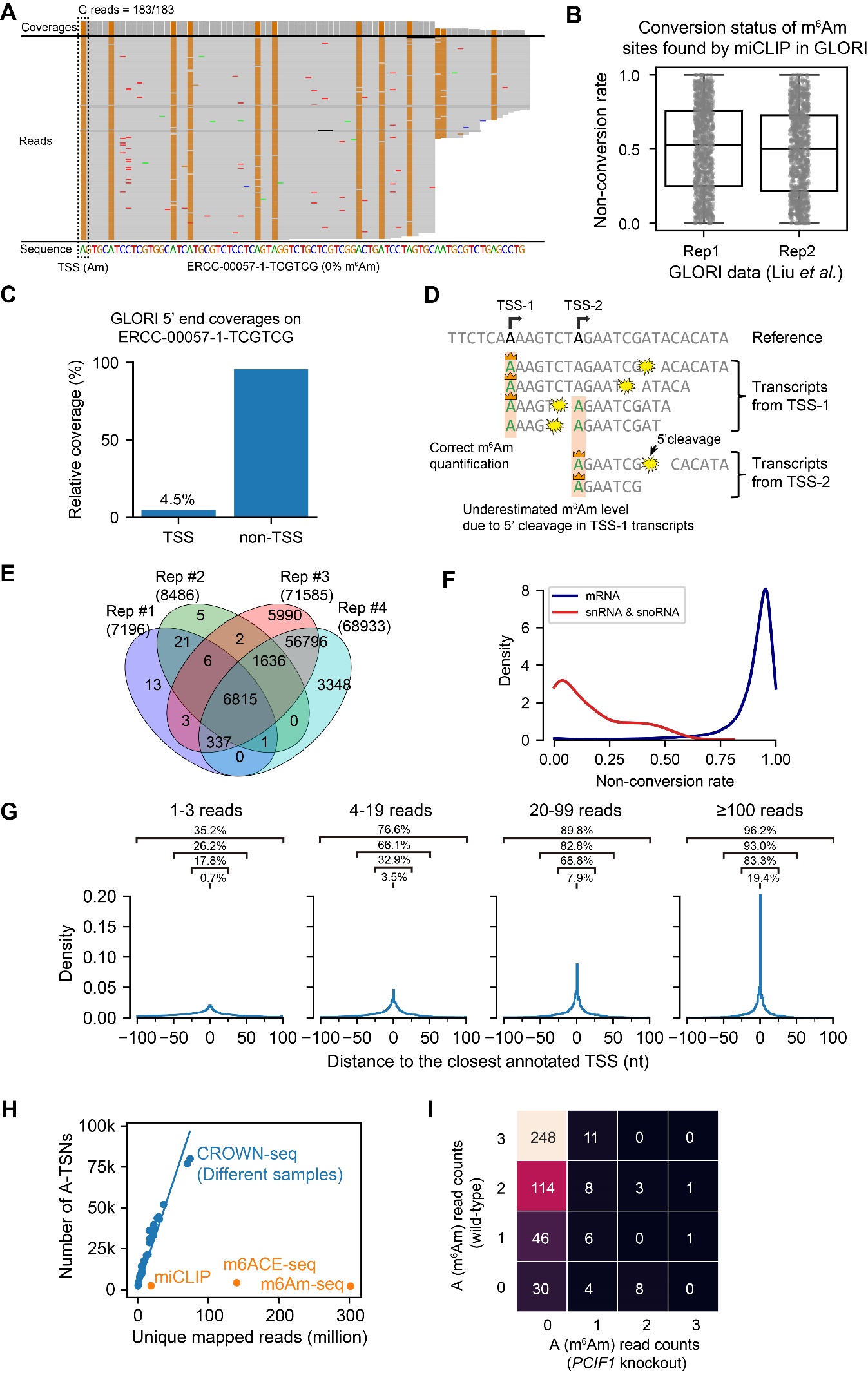

**Figure S2. Corresponding to Figure 2.**

**(A)** GLORI can completely convert 5’ end Am in a transcript synthesized by *in vitro* transcription. Shown is the IGV plot demonstrating the coverage and individual reads mapped to the TSS of ERCC-00057-1-TCGTCG transcript. The reference sequence is also shown at the bottom. In this plot, the bases that match the reference are shown in gray, while G mismatches are shown in brown. The A-to-G mutation in the TSS position indicates successful A-to-I conversion during sodium nitrite treatment. In this assay, the 5’ Am in ERCC-00057-1-TCGTCG transcript was made by using m7GpppAmG analog during *in vitro* transcription (see **Methods**). The ERCC-00057-1-TCGTCG transcript was decapped and spiked into the RNA sample for GLORI experiment.

**(B)** GLORI underestimated m^6^Am stoichiometry in m^6^Am sites found by miCLIP. In this analysis, we reanalyzed the reads in HEK293T GLORI libraries (GSE210563) generated by Liu et al. ^8^. To obtain the non-conversion rate of m^6^Am sites, we extracted the 5’ ends mapped to miCLIP sites found by Boulias *et al.* ^5^. 1084 and 1067 m^6^Am sites were analyzed in each replicate, respectively.

**(C)** Only a few reads in GLORI can mapped to the desired TSS. In this analysis, we analyzed the GLORI mapping result of ERCC-00057-1-TCGTCG transcript in **(A)**. We calculated the relative ratio of reads whose 5’ end mapped to not failed to map to the desired TSS of the ERCC-00057-1-TCGTCG. Only ~4.5% of the reads can be correctly mapped to the TSS.

**(D)** A schema showing the reason why m^6^Am stoichiometry is incorrectly estimated in the transcriptome. In transcriptome, a gene can have multiple TSSs, whose 5’ ends can overlap with each other. As a result, the A-TSN from TSS-2 can also be transcribed as an internal A base in transcripts generated by TSS-1. During the GLORI experiment, 5’ cleavage happens very frequently, which results in massive 5’ ends from internal A’s. Since the GLORI library is unable to distinguish between the “true” 5’ ends with m^7^G cap and the “false” 5’ ends with monophosphate, both “true” 5’ ends and “false” 5’ ends are counted for TSS-2. Because most of the internal A’s are not N6-methylated, the “false” 5’ ends will dilute the m^6^Am signal from the “true” 5’ ends and result in underestimated m^6^Am stoichiometry.

**(E)** CROWN-seq shows high reproducibility in m^6^Am identification. In the Venn diagram, four replicates of HEK293T CROWN-seq results were compared. Among the replicates, replicates #1 and #2 are two technical replicates of the same biological sample; while replicates #3 and #4 are the two technical replicates of another biological sample. In this analysis, A-TSNs with at least 20 reads mapped were included. Since replicate #1 and #2 have lower sequencing depth (~3 million reads each) than replicate #3 and #4 (~70 million reads each), the numbers of A-TSNs reported by replicate #1 and #2 are much lower than that by replicate #3 and #4.

**(F)** Kernel density estimate (KDE) plot for the distribution of non-conversion rates of A-TSNs belonging to mRNA and snRNA/snoRNA reported by CROWN-seq. In this analysis, all sequencing results from HEK293T replicates were merged. Only A-TSNs with at least 20 reads were analyzed.

**(G)** The distance between CROWN-seq identified TSSs and annotated TSSs in Gencode v45. In this plot, the TSSs identified in CROWN-seq replicate 3 were analyzed. 2,054,368, 333,959, 120,378, and 63,849 TSSs with 1-3, 3-19, 19-99, and ≥100 mapped reads were shown respectively. The percentage of TSSs well overlapped with annotation, as well as TSS located within the [-25, 25], [-50, 50], and [-100, 100] regions proximal to the annotated TSSs were also indicated.

**(H)** Corresponding to **Figure 2J**, shown are the number of uniquely mapped reads (X-axis) and the number of called m^6^Am or A-TSN in different methods. For CROWN-seq, all libraries used in this study are shown in dots. We performed linear regression (shown in line) to calculate the sensitivity of CROWN-seq (i.e., the slope of linear regression) which is shown in **Figure 2J**.

**(I)** CROWN-seq exhibited high TSS mapping accuracy in low-coverage sites. In this heatmap, each block represents A-TSNs with a certain number of A reads (i.e., m^6^Am reads) mapped in wild-type (Y-axis) and *PCIF1* knockout (X-axis) HEK293T. Only A-TSNs have 3 reads mapped in wild-type and 3-39 mapped in *PCIF1* knockout are shown. For example, shown in the upper left corner, there are 248 A-TSNs that have 3 A reads in wild-type (i.e., 100% non-conversion) and 0 A reads (i.e., 0% non-conversion) in *PCIF1* knockout cells.

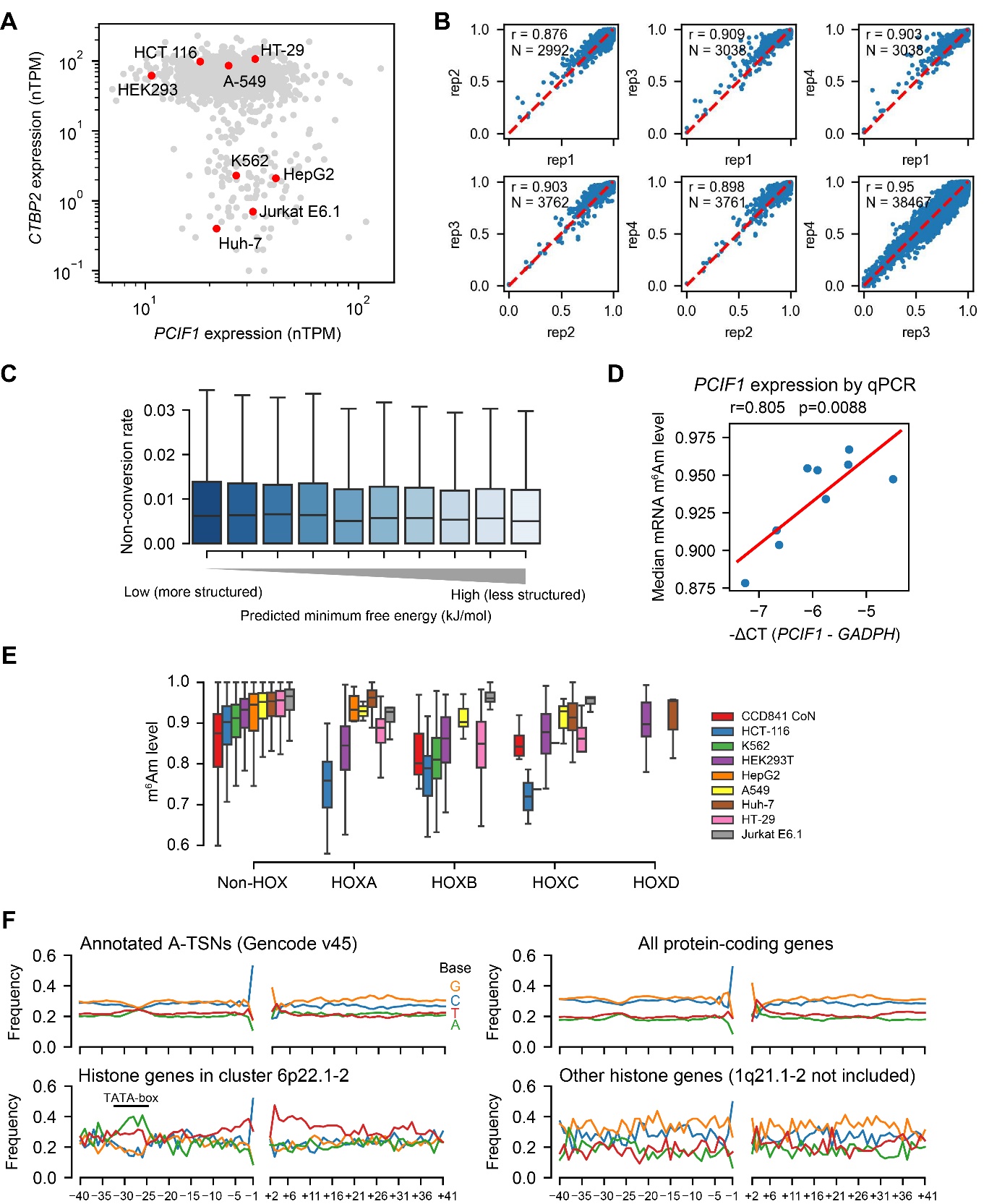

**Figure S3. Corresponding to Figure 3.**

**(A)** Several cell lines exhibit low *CTBP2* expression. Shown are normalized RNA expression results from the Human Protein Atlas (<https://www.proteinatlas.org/download/rna_celline.tsv.zip>)^9^.

**(B)** The 50 reads threshold yields highly correlated non-conversion rates among replicates. Shown are the pairwise comparisons of non-conversion rates of A-TSNs quantified in different HEK293T replicates. In this analysis, A-TSNs were required to have at least 50 reads mapped. As a result, the A-TSNs in different replicates have median read counts of 105-149. High correlations (Pearson’s r) in non-conversion rates were found between replicates.

**(C)** RNA secondary structure is unlikely to affect m^6^Am quantification accuracy in CROWN-seq. Shown are the non-conversion rates of A-TSNs grouped by the degrees of RNA secondary structure in *PCIF1* knockout HEK293T cells, which have no m^6^Am in mRNA. To obtain the degree of RNA secondary structure of each 5’ end, we calculated the minimum free energy of A-TSN plus the first 30 nt downstream sequence by RNAfold^10^. 18,235 A-TSNs were binned into 10 bins based on the quantile of minimum free energy. For each bin (from left to right), the medium minimum free energies are -11.4, -9.2, -7.9, -6.9, -6.0, -5.3, -4.5, -3.6, -2.6, and -0.9 kJ/mol.

**(D)** Comparing median mRNA m^6^Am stoichiometry and *PCIF1* expression estimated by RT-qPCR. In this assay, RT-qPCR on PCIF1 and GADPH transcripts was performed among all nine different cell lines. The relative expression levels of PCIF1 were normalized by the expression of *GAPDH*. Linear regression was performed to obtain Pearson’s r and P-value.

**(E)** Boxplots showing the m^6^Am levels of HOX genes among different cell lines.

**(F)** Histone genes in cluster 6p22.1-2 exhibited distinct sequence context in the core promoter. Shown are the flanking sequences of the 146,285 A-TSN that annotated in Gencode v45 (upper left), 104,733 A-TSNs mapped to protein-coding genes in CROWN-seq (upper right), 228 histone genes in cluster 6p22.1-2 (lower left), and 119 histone genes not in neither cluster 6p22.1-2 nor 1q21.1-2 (lower left). Since a few (14) A-TSNs were found in cluster 1q21.1-2, A-TSNs in cluster 1q21.1-2 are not shown.

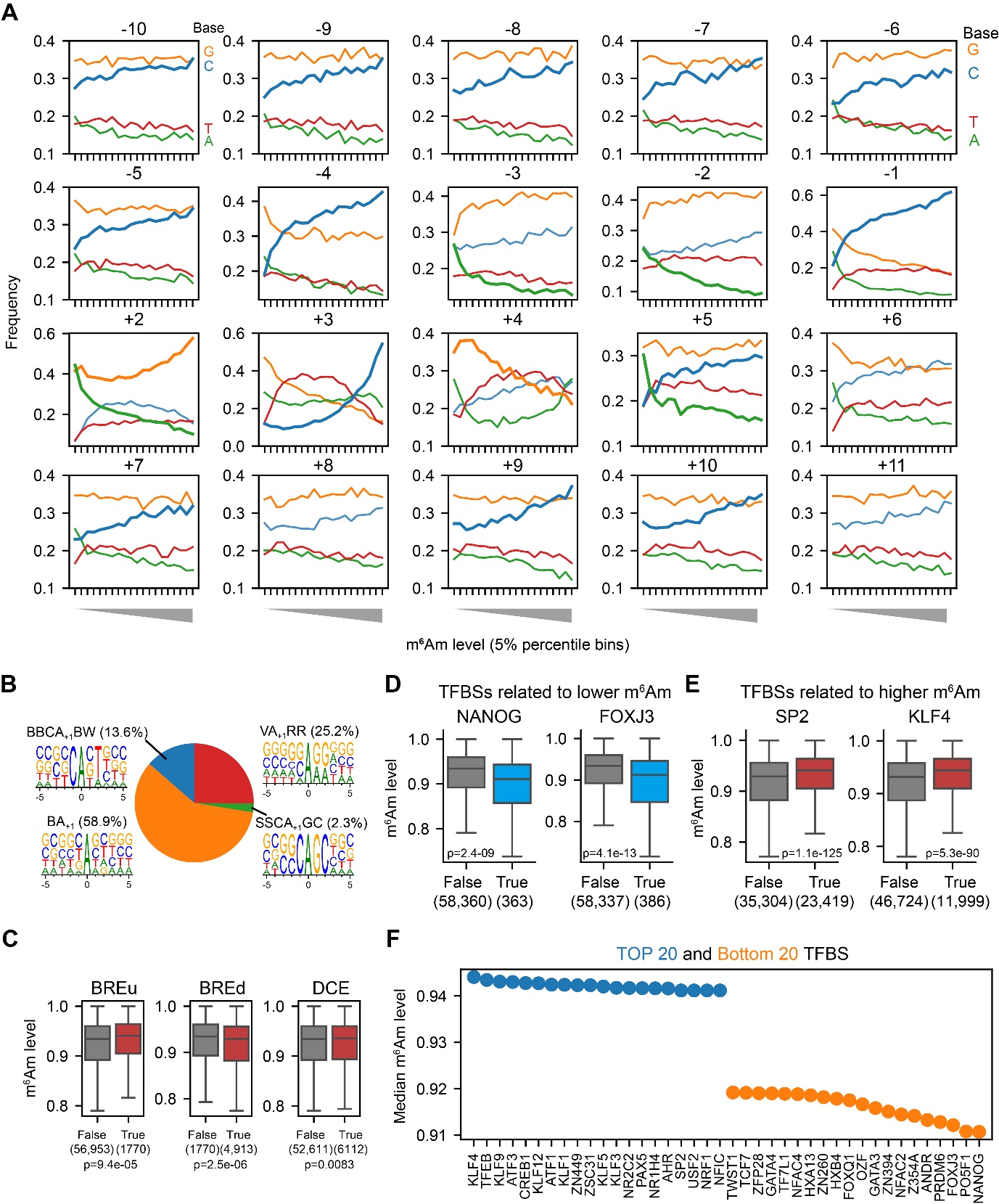

**Figure S4. Corresponding to Figure 4.**

**(A)** Corresponding to **Figure 4A**, the base preference in -10 to +11 core promoter region of the 58,723 A-TSNs in HEK293T cell. To examine the correlation between m^6^Am stoichiometry and base compositions, we equally binned the A-TSNs by m^6^Am stoichiometry into twenty 5-percentile bins. For each bin, the frequency of A, T, C, and G bases are shown.

**(B)** A pie plot showing the fractions of A-TSNs using different TSS motifs. In this plot, 58,723 A-TSNs in HEK293T cells are shown, where 1,376 are from the SSCA_+1_ GC motif, 7,981 are from BBCA_+1_BW motif, 14,788 are from VA_+1_RR motif, and 34,578 are from BA_+1_ motif.

**(C)** Examining the relationship between core promoter elements and m^6^Am stoichiometry. In this analysis, core promoter elements (BREu, BREd, and DCE) were defined according to the description by Haberle and Stark ^11^ (indicated as “True”). BREu-containing promoters were defined as promoters that contain SSRCGCC (S=C/G, R=A/G) at -38 to -32; BREd-containing promoters were promoters that contain RTDKKKK (D=A/G/T, K=G/T) at -23 to -17; DCE containing promoters were defined as promoters which contain CTTC at +6 to +11, or CTGT at +16 to +21, or AGC at +30 to +34. Among these elements, BREu and BREd are related to TFIIB binding, while DCE is related to TAF1 binding^11^.

**(D)** and **(E)**, Examples of m^6^Am stoichiometry for A-TSNs containing specific transcription factor-binding sites. In this analysis, m^6^Am stoichiometry of the 58,723 A-TSNs in HEK293T was used; transcription factor-binding sites within the -50 to +51 region were identified by FIMO^12^ based on the HOCOMOCO v11 core motifs database^13^. Transcripts containing NANOG and FOXJ3-binding sites (indicated as “True”) exhibit relatively lower m^6^Am stoichiometry, while SP2 and KLF4-binding sites are associated with higher relative m^6^Am stoichiometry. The overall effect size is relatively small, which suggests that transcription factors may not have large effects on m^6^Am stoichiometry. P-values, Student’s t-test, two-sided.

**(F)** Some transcription factor binding sites (TFBS) are related to higher or lower m^6^Am stoichiometry. In this figure, the top 20 and bottom 20 m^6^Am-related TFBS are shown. The m^6^Am stoichiometry and the -50 to +51 sequence of the 58,723 A-TSNs in HEK293T cells were analyzed. FIMO^12^ was used to scan for core motifs in the HOCOMOCO v11 database. Only TFBS occurred in the flanking sequence of at least 200 A-TSNs are shown.

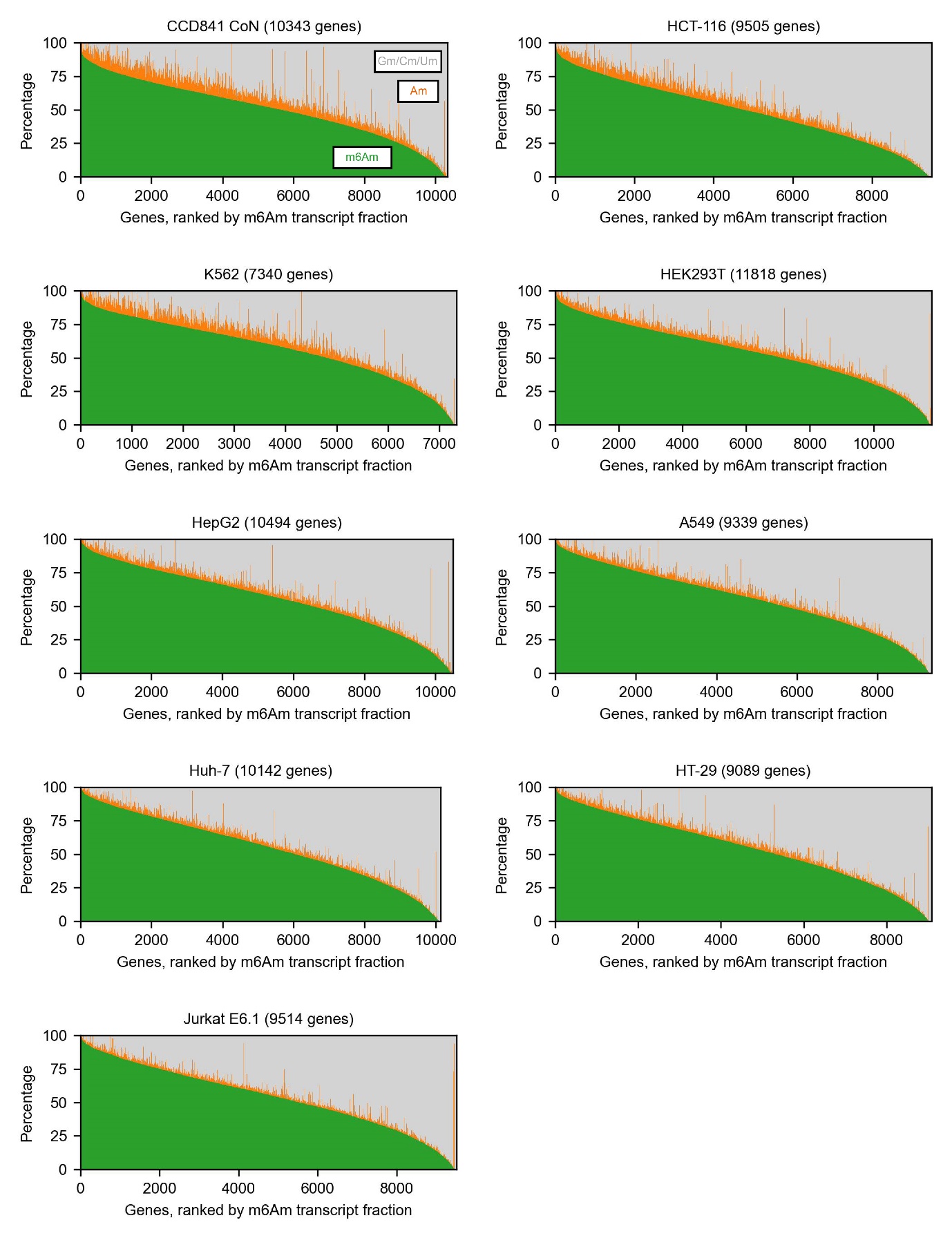

**Figure S5. The fraction of different transcription-start nucleotides of a gene among different cell lines.**

Similar to **Figure 1D**, the fractions of transcripts starting with m^6^Am, Am, and other bases in different cell lines. This estimation is based on CROWN-seq. Only genes with at least 50 mapped reads were shown.

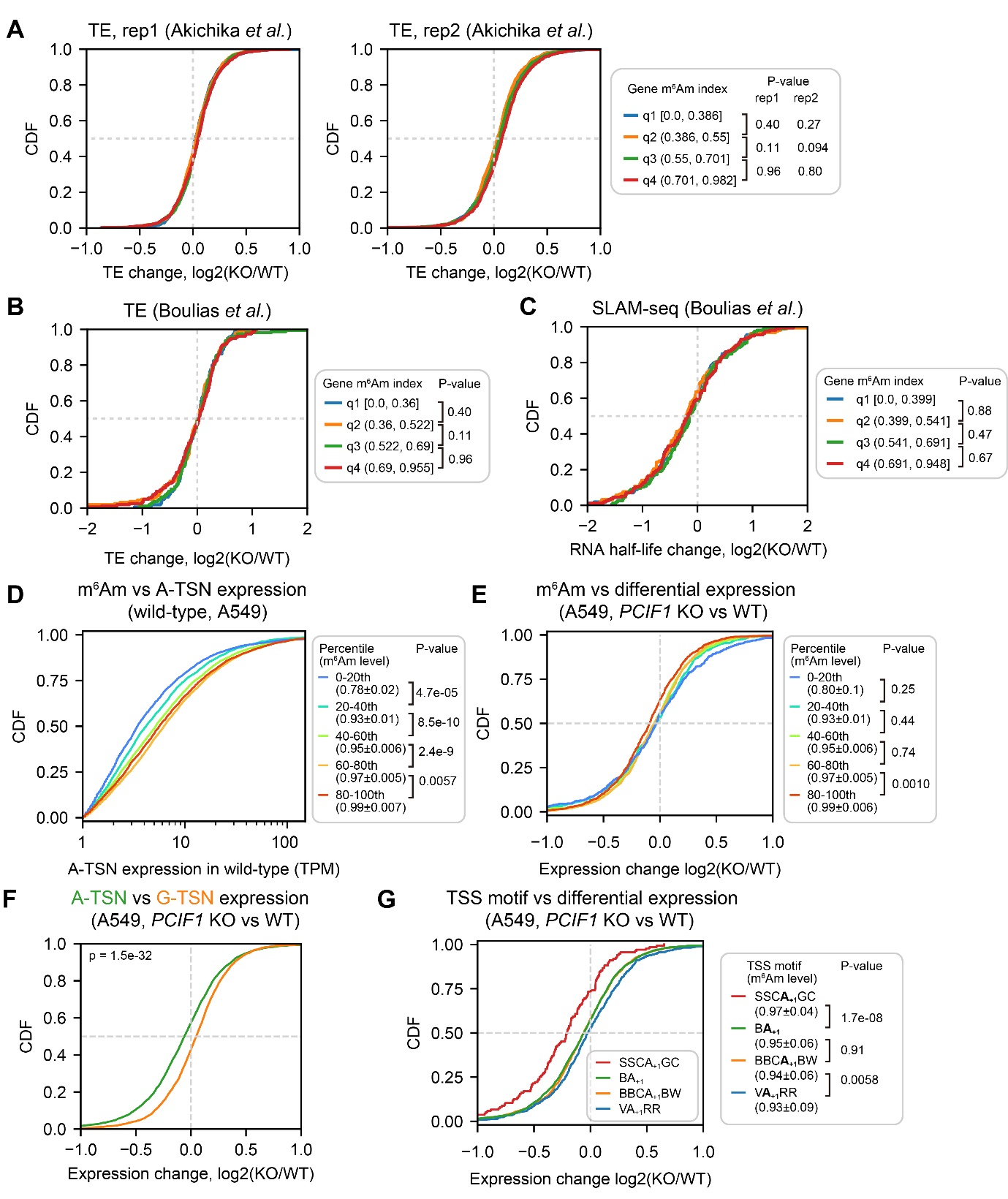

**Figure S6. Corresponding to Figure 5.**

**(A)** *PCIF1* knockout did not alter translation efficiency in HEK293T cells. Shown are ribosome profiling results by Akichika *et al.* ^4^. In this analysis, 3,325 genes were equally binned into quantiles by gene m^6^Am index. P-values, Student’s t-test, two-sided.

**(B)** *PCIF1* knockout did not alter translation efficiency in HEK293T cells. Shown are ribosome profiling results by Boulias *et al.* ^5^. In this analysis, 637 genes were equally binned into quantiles by gene m^6^Am index. P-values, Student’s t-test, two-sided.

**(C)** *PCIF1* knockout did not alter RNA stability in HEK293T cells. Shown are RNA half-life results estimated by SLAM-seq by Boulias *et al.* ^5^. In this analysis, 666 genes were equally binned into quantiles by gene m^6^Am index. P-values, Student’s t-test, two-sided.

**(D)** m^6^Am stoichiometry is positively related to A-TSN expression in wild-type A549 cells. In this cumulative distribution plot, A-TSNs with expression quantified by ReCappable-seq in different m^6^Am stoichiometry bins are shown. To group different A-TSNs, we first binned 58,723 A-TSNs into five bins based on m^6^Am stoichiometry quantified by CROWN-seq. Then the A-TSNs detected in ReCappable-seq were annotated by the predefined bins. In total, 5,125, 6,962, 7,991, 8,368, and 8,009 A-TSNs are shown in each bin (from low m^6^Am to high m^6^Am). These A-TSNs have an average TPM ≥ 1 in two ReCappable-seq replicates and coverage ≥50 in CROWN-seq. P-values, Student’s t-test for TPM (log-transformed), two-sided.

**(E)** A-TSNs in high m^6^Am stoichiometry are more susceptible to *PCIF1* knockout. Shown are the cumulative distributions of A-TSN expression change in A549 cells upon *PCIF1* knockout. The differential expression of A-TSN was calculated by DESeq2^14^. Similar to **(D)**, the A-TSNs were binned based on the m^6^Am stoichiometry. In total, 481, 738, 973, 1,212, and 1,087 A-TSNs are shown in each bin (from low m^6^Am to high m^6^Am). These A-TSNs have baseMean (i.e., the average of the normalized count among replicates) ≥100 during the differential expression test (2 replicates for both wild-type and *PCIF1* knockout) and coverage ≥50 in CROWN-seq. P-values, Student’s t-test, two-sided.

**(F)** Shown are cumulative distributions of expression changes of A-TSNs and G-TSNs. 4,975 A-TSNs and 3,510 G-TSNs with expression levels quantified by ReCappable-seq are shown. These TSNs have an average baseMean ≥100 of two replicates. P-values, Student’s t-test, two-sided.

**(G)** Similar to **(F)**, A-TSNs using different TSS motifs exhibited different changes in expression upon *PCIF1* knockout. In total, 135 A-TSNs using SSCA_+1_GC, 2.502 A-TSNs using BA_+1_, 1,131 A-TSNs using BBCA_+1_BW, and 723 A-TSNs using VA_+1_RR are shown. These A-TSNs have baseMean ≥100 during the differential expression test (2 replicates for both wild-type and *PCIF1* knockout) and coverage ≥50 in CROWN-seq. P-values, Student’s t-test, two-sided.

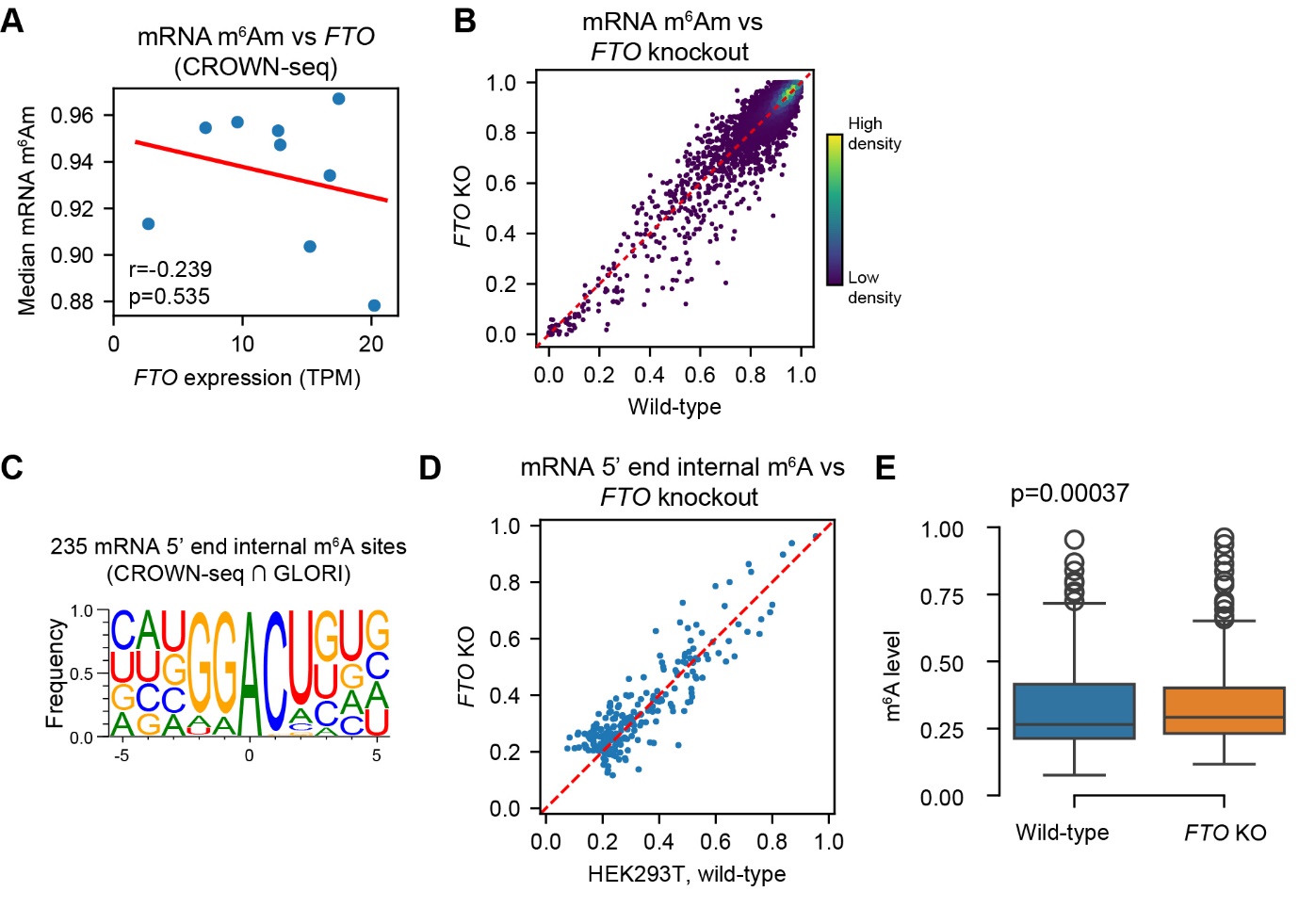

**Figure S7. Corresponding to Figure 6.**

**(A)** *FTO* expression correlates weakly to mRNA m^6^Am stoichiometry. Shown are *FTO* expression measured by CROWN-seq (X-axis) and median mRNA m^6^Am levels among nine different cell lines.

**(B)** FTO depletion causes small changes in mRNA m^6^Am stoichiometry. In this scatter plot, only mRNA A-TSNs which have at least 50 reads in both wild-type and *FTO* knockout cells were shown. m^6^Am levels were estimated by CROWN-seq.

**(C)** The internal m^6^A sites identified by both CROWN-seq and GLORI^8^ showed a classic DRACU motif.

**(D)** and **(E)** *FTO* knockout showed a subtle effect in changing the stoichiometry of internal m^6^A around 5’ ends. Shown are **(D)** a scatter plot comparing m^6^A stoichiometry between wild-type and *FTO* knockout cells, and **(E)** a boxplot showing the overall m^6^A stoichiometry difference. In **(E)**, Student’s t-test (two-sided) was performed to calculate the significance of the difference in m^6^A stoichiometry.

**Note S1. The comparison of TSS mapping methods.**

Currently, there are several types of TSS mapping methods. CAGE^15^ and TSS-seq^16^ are the two most popular methods being used.

CAGE was tested to have the highest precision and sensitivity over other TSS mapping methods^17^, except for ReCappable-seq. However, CAGE has two limitations. First, CAGE relies on template switching. Template switching is a process in that reverse transcriptase can “jump” onto a template switching oligo, which contains an adapter sequence when the reverse transcriptase reaches the end of the RNA template. Template switching is very convenient in producing full-length cDNA without ligating adapter. However, template switching is not precise for transcription-start nucleotide identification, because template switching can introduce non-template bases (normally C’s) into the cDNA between the template and the adapter. It is very difficult to completely remove the non-template bases in the CAGE library because the number of incorporated non-template bases is uncertain^18^. As a result, compared with TSS-seq and ReCappable-seq, CAGE can mistakenly assign TSSs within the same CAGE peak (see **Figure S1 A, B**). Second, the most widely used CAGE protocol^15^ contains an oxidation step, which results in massive indels and mutations in the cDNA. These indels and mutations can result in inaccurate alignments. Third, “strand invasion” can cause TSS artifacts in CAGE^18^. Strand invasion is the process that reverse transcriptase mistakenly terminates and switches onto the template switching oligo before reaching the end of a template. Strand invasion can result in false positive TSSs in internal RNA positions.

TSS-seq is another available method in TSS mapping. TSS-seq relies on several enzymatic steps to remove non-m^7^G-capped 5’ end backgrounds in the sample. After removing undesired 5’ ends, the m^7^G cap is released and a 5’ adapter is ligated to the RNA 5’ ends. In theory, this procedure can result in precise 5’ end maps. However, tested by Adiconis *et al.* ^17^, TSS-seq exhibited low precision, sensitivity, and accuracy in TSS mapping. The low performance of TSS-seq is due to the incomplete removal of the non-m^7^G-capped 5’ end backgrounds.

ReCappable-seq^19^ can be considered as an improved TSS-seq. Recappable-seq overcomes the 5’ end background clean-up issue. In ReCappable-seq, the m^7^G caps of RNA polymerase II transcribed RNA is replaced by 3´-Desthiobiotin-G caps. The recapped RNAs can thus be enriched on streptavidin beads. During high-stringency washing, the 5’ end background can be completely removed. Thus, ReCappable-seq exhibited extremely high specificity in mapping transcription-start nucleotides.

**Note S2. Selection of m^6^Am identification cutoffs.**

The choice of parameters can significantly affect the accuracy of TSS maps and the precision in m^6^Am quantification. In this study, we used several different parameters in defining TSS signals from ReCappable-seq and CROWN-seq.

In **Figure 1** and **S1**, for the preliminary analyses with ReCappable-seq, we defined TSSs as those with ≥1 TPM coverage as previously used^19^. Notably, this threshold is empirical and subjective for TSS identification. This threshold can result in false negatives, especially for those TSS with expression levels a bit lower than 1 TPM.

In **Figure 2** and **S2**, to define A-TSNs in CROWN-seq, we first called high-confidence A-TSNs which at least mapped by 20 reads. This threshold was used in a previous m^5^C mapping analysis^20^. The ≥20 reads threshold can yield acceptable precision in m^6^Am stoichiometry estimation. When an A-TSN is mapped by 20 reads, the quantification precision is 0.05 (1/20). With this threshold, the median coverage of the A-TSNs is ~40-60 among samples, which means precision at 0.017-0.025. Notably, according to the analysis shown in **Figure 2H** and **Figure S2I**, CROWN-seq exhibited very high accuracy in TSN mapping even for the TSNs mapped by 3 reads. Although this threshold allows us to roughly estimate m^6^Am stoichiometry, the variability of the quantified stoichiometry can be high when the read depth is low (particularly for A-TSNs with <50 reads). Thus, we used another criterion while generating the m^6^Am landscape among different cell lines.

For m^6^Am landscape profiling (**Figure 3-6, S3-S6**), we want to precisely compare the m^6^Am stoichiometry between different cell lines. We first merged all the reads from different biological and/or technical replicates to obtain higher read depths for each cell line. We then increased the threshold of sequencing depth so that only A-TSNs mapped by ≥50 reads were quantified. With this threshold, the minimum precision is set to 0.02 (1/50). In practice, this threshold results in medium read coverage at ~130-150 reads, which indicates precision at 0.0067-0.0077. The high coverage also results in low variability in m^6^Am quantification between replicates (see **Figure S3B**).
